# Spatial transcriptomics for gene discovery identifies *Slc13a5* as a modulator of bone mechanoadaptation

**DOI:** 10.64898/2026.03.11.711126

**Authors:** Quentin A. Meslier, Alec T. Beeve, Arushi Gupta, Daniel Palomo, Samia Saleem, Serena Eck, Lisa Lawson, John Shuster, Michelle Brennan, Naomi Dirckx, Matthew J. Silva, Erica L. Scheller

## Abstract

Bone is a dynamic tissue that continuously adapts its structure in response to mechanical loading, an essential process for maintaining skeletal health. However, this adaptive capacity declines with aging, contributing to increased fragility and fracture risk. Developing therapeutic strategies that preserve or restore bone mechanoadaptation in patients with increased bone fragility requires identifying key molecular regulators of this process. We applied spatial transcriptomics (GeoMx, NanoString) to characterize gene expression changes induced by mechanical loading in the murine tibia, focusing on periosteal and bone compartments in regions under tension and compression. Spatial data were validated and cross-compared with previously published bulk RNA-seq and laser-capture microdissection datasets, identifying a set of 12 genes consistently regulated by loading across independent platforms and laboratories. As part of a functional analysis, we selected *Slc13a5*, a citrate transporter implicated in bone mineralization and metabolism. Conditional deletion of *Slc13a5* in osteolineage cells using Osteocalcin-Cre significantly increased the loading-induced mineralizing surface in tensile regions compared with Cre⁻ *Slc13a5^fl/fl^* littermates. In addition, *Slc13a5* cKO mice exhibited lower resorption around the neutral axis after loading compared to controls. Together, these findings identify *Slc13a5* as a regulator of bone adaptation in regions experiencing low mechanical stimulation and suggest it as a potential therapeutic target for conditions characterized by impaired mechanoadaptive responses. This study highlights spatial transcriptomics as a powerful gene discovery framework for bone, enabling identification of novel targets to understand mechanisms and develop therapies.

**Graphical Abstract:** 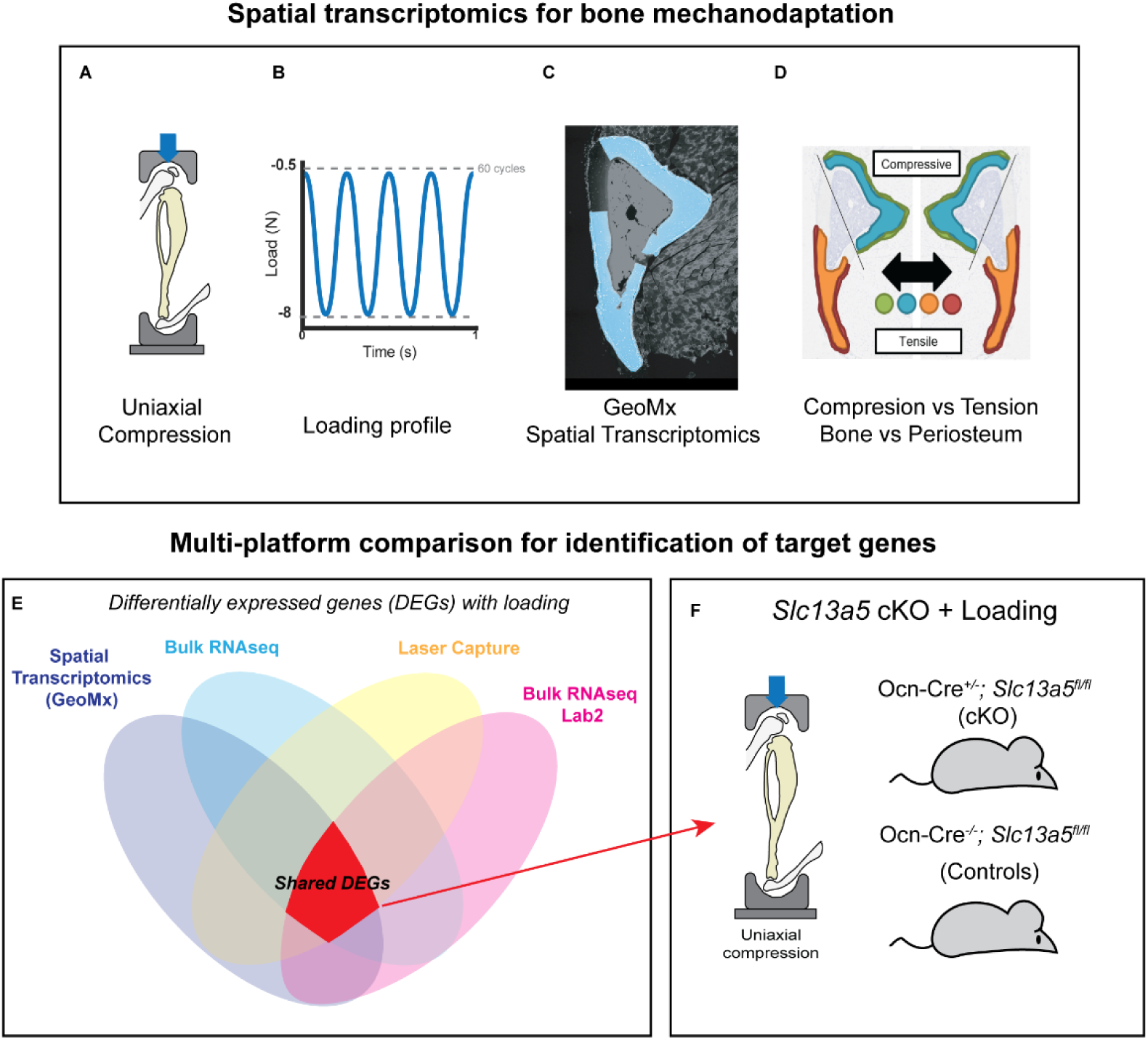

## Introduction

Bone is a dynamic tissue that adapts its structure and mass in response to mechanical loading. Mechanoadaptation is essential for maintaining skeletal integrity and is disrupted with aging [1], [2], [3]. Although the anabolic effects of mechanical loading are well established, the molecular mechanisms linking local mechanical stimuli to spatially patterned bone formation remain incompletely understood. Specifically, how distinct bone subregions respond transcriptionally to different mechanical environments remains a key unanswered question.

Previous studies have used targeted approaches such as RT-qPCR, in situ hybridization, and immunohistochemistry [2], [4] to examine candidate mechanosensitive pathways, while bulk RNA sequencing has revealed broader transcriptional responses to loading [5], [6]. These studies identified pathways involved in osteogenesis, extracellular matrix organization, and mechanotransduction. However, bulk approaches lack spatial resolution and cannot capture heterogeneity in mechanical stimuli across the bone cross-section or along its length. In the uniaxial tibial loading model [7], the posterior-lateral surface is under compression and exhibits the greatest bone formation, whereas the anterior-medial surface is under tension and shows a more limited anabolic response [2]. This spatial distribution arises from the natural curvature and geometry of the tibia. The reduced anabolic response on the tensile surface has been attributed to lower mechanical stimulus magnitude during axial loading.

Capturing gene expression within these mechanically distinct microenvironments is critical for understanding site-specific bone adaptation. Laser capture microdissection [8] provides compartment-level insight into periosteal versus cortical bone responses but has poor efficiency and has limited spatial precision. Whole-mount labeling enables visualization of gene expression along the bone length but is restricted to a small number of predefined targets per run [9]. Spatial transcriptomic approaches overcome these limitations by enabling spatially resolved, full-transcriptome analysis within intact tissue sections [10].

In this study, we applied the GeoMx Digital Spatial Profiler (DSP) to compare loading-induced gene expression changes in defined bone subregions of the mouse tibia, including periosteum versus cortical bone and compressive versus tensile regions. We implemented a three-level analytical framework to validate the platform against bulk RNA-seq and laser capture datasets and leveraged its spatial resolution to uncover region-specific mechanoadaptive transcriptional programs. Finally, we used this approach to identify and functionally interrogate a novel candidate regulator of bone mechanoadaptation: solute carrier family 13 member 5 (*Slc13a5)*. *Slc13a5* is a citrate transporter whose deletion can improve whole-body metabolic homeostasis, protecting aged or high-fat-diet–fed mice from adiposity and insulin resistance [11]. Citrate is highly abundant in bone, which serves as the body’s primary reservoir of this metabolite. Within bone, citrate becomes incorporated into the hydroxyapatite-collagen composite of the mineralized matrix and also supports cellular metabolism as a mitochondrial substrate [12], [13]. *Slc13a5* mediates citrate uptake and has been reported to be expressed in late osteoblasts and osteocytes [12]. Thus, *Slc13a5* represents a promising and largely unexplored therapeutic target for modulating bone mechanoadaptation.

Together, this work provides a spatially resolved molecular framework for bone mechanoadaptation and highlights new genes and pathways that may be targeted to modulate skeletal responses to loading.

## Materials and Methods

### Animals

This study received full approval from the Washington University Institutional Animal Care and Use Committee (IACUC). C57BL/6J female mice were obtained from Jackson Laboratories (Strain #000664). A bone-specific *Slc13a5* conditional knockout (cKO) model was generated by crossing osteocalcin *Ocn*-Cre mice with *Slc13a5* floxed (*Slc13a5^fl/fl^*) mice [12]. The resulting progeny included *Slc13a5* conditional knockout female mice (*Ocn*-Cre^+/−^; *Slc13a5^fl/fl^*, hereafter referred to as *Slc13a5^cKO^*) and female littermate controls (*Ocn*-Cre^−/−^; *Slc13a5^fl/fl^*). Mice were housed in group cages within the Washington University animal facility, maintained at a temperature of 22-25°C, following a 12-hour light and 12-hour dark cycle with continuous access to water and chow (Purina 5053, Saint Louis, MO, USA). Mice were used for unilateral tibial loading protocols at 5 months of age.

### Unilateral in vivo tibial compression

Mice were anesthetized using 2-3% isoflurane. The right tibia was secured at the knee and heel in custom fixture of a material testing system (Instron Dynamite 8841). Uniaxial compressive loading was applied cyclically to the right tibia once daily for 5 days Monday to Friday (M-F) for spatial transcriptomics (n=3); or 5 days a week for 2 weeks (M-F) for loading-induced bone accrual in *Slc13a5^cKO^* (n=9) and *Slc13a5^fl/fl^* (n=8). A loading bout consisted of 60 cycles at 4 Hz with a peak compressive strain of -2200 µε, along with a concurrent peak tensile strain of 1200 µε, [2], [14], [15]. Based on prior strain gauge analysis [15], the force necessary to produce peak compressive strains of −2200 µε was estimated to be −8N. A preload of -0.5N was applied to keep the tibia in place. Mice received 0.05-0.1 mg/kg buprenorphine subcutaneously following each loading session for analgesia. The loading experiment is summarized in Fig.1.

**Figure 1:**
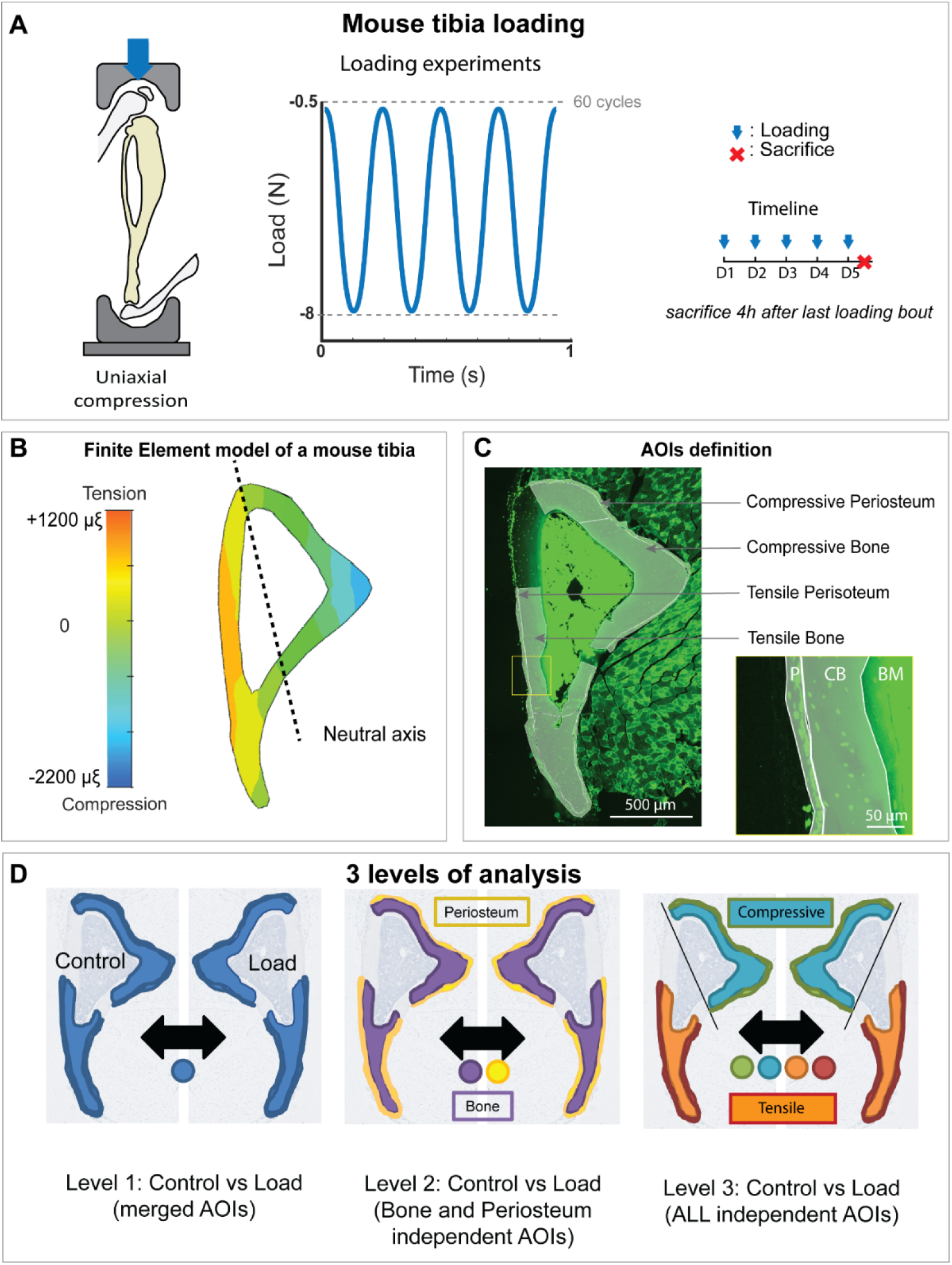
Loading experiment summary. A) Right tibiae were loaded using a uniaxial compression model. Cyclic compression of -8N was applied for 5 days. B) Finite element model of a mouse tibia showing typical mechanical stimulus pattern in the tibia midshaft under uniaxial compression. The posterior-lateral side of the tibia is in compression whereas the anterior-medial side is in tension (Adapted from Meslier et al. 2022 [22]). C) Mouse tibia section used for spatial transcriptomics showing the AOI definition in the GeoMx software. P= Periosteum. CB= Cortical Bone. BM = Bone Marrow. D) Illustration showing the combination of the AOIs for the 3 different level of analysis.

### Tissue preparation and sectioning

Loaded and control legs for spatial transcriptomics were harvested 4 hours after the final loading bout following euthanasia by CO₂ inhalation. Benchtops and tools were pre-cleaned with RNase Away. Bilateral tibiae with intact periosteum were trimmed to remove excess muscle and placed in 15 mL conical tubes containing 10% neutral buffered formalin for 24 h at room temperature with gentle agitation. Fixed tissues were washed 3 × 30 min in 1× PBS in the same tubes. Decalcification was performed in 14% EDTA (pH 7.2), refreshed every 2–3 days for 14 days. Bones were then washed 3 × 30 min in Milli-Q water and dehydrated through 30%, 50%, and 70% ethanol (3 × 30 min each) before submission to the Musculoskeletal Histology and Morphometry Core (Washington University, St. Louis, USA) for paraffin embedding. Blocks were stored at 4°C until use. Transverse sections (5 µm thick) were collected ∼5 mm distal to the tibial plateau and mounted on coated glass slides (Agilent Dako IHC FLEX, K8020) for use within one week. Loaded and control legs from *Slc13a5*^cKO^ and *Slc13a5^fl/fl^* mice were collected 18 days after loading initiation (7 days after the final loading bout). Muscles and surrounding soft tissues were removed, bones were fixed overnight in 10% neutral buffered formalin at room temperature, dehydrated through graded ethanol (20%, 50%, 70%), and stored in 70% ethanol at 4°C.

### GeoMx pipeline for spatial capture and analysis of regional gene expression

RNAscope was used to verify RNA preservation of the samples before proceeding to spatial transcriptomics as described previously [16]. The detailed protocol is available in the supplemental methods. Spatial transcriptomic profiling was then performed using the NanoString GeoMx® DSP platform, which preserves the spatial organization of gene expression within tissue sections. The GeoMx DSP platform enables whole-transcriptome analysis of mouse tissues using barcoded, photocleavable probes to capture spatially resolved gene expression. Prepared sections were mounted on GeoMx-compatible slides (Agilent Dako, IHC FLEX, K8020), and probes targeting selected transcripts were hybridized to accessible RNA. Bone samples required optimization of pretreatment steps to improve section adherence and antigen retrieval. Briefly, slides were dried overnight at 60°C. Samples were then deparaffinized, rehydrated, and pretreated with hydrogen peroxide. Slides were incubated in MilliQ water heated at 75°C followed by a pepsin treatment to complete the antigen retrieval. GeoMx probes were hybridized overnight at 40°C prior to continuing with kit-recommended washes and counterstain (Supplemental methods).

Areas of interest (AOIs) were manually defined under microscopic visualization using the GeoMx instrument imaging interface, guided by nuclear counterstain (Fig.1C). Regions experiencing maximal mechanical stimulus (Fig.1B) during uniaxial tibial compression were targeted, including compressive bone and periosteum (posterior–lateral side) and tensile bone and periosteum (anterior–medial side) (Fig.1C-D). The GeoMx instrument selectively illuminated each AOI to release hybridized probes, which were collected into separate wells while preserving spatial correspondence. GeoMx libraries were pooled and sequenced on an Illumina NovaSeq 6000. Raw FASTQ files were processed using the GeoMx NGS Pipeline to generate DCC files (counts per Readout Tag Sequence ID [RTS-ID]), including read trimming, alignment, and PCR duplicate removal. Data were analyzed using the GeoMx Tools R package, enabling identification of region-specific transcriptional signatures (Fig.1D).

For quality control, default NanoString-recommended parameters were used. To balance stringency and data retention, the minNegativeCount cutoff was lowered and the minArea threshold increased, preserving biologically relevant features while removing artifacts. Dimensionality metrics were monitored across QC steps to detect feature loss and prevent unnecessary data exclusion. Segment-level quality control (QC) ensured sequencing quality and adequate tissue sampling, followed by bioprobe-level QC to identify and remove globally underperforming probes. A gene-level count matrix was then generated, and the limit of quantification (LOQ) was calculated for each segment to filter genes with low signal. Data were normalized using third quartile (Q3) normalization, and unsupervised dimensionality reduction was applied to cluster ROIs based on global gene expression. The full dataset is available at [GSE324027]. These QC steps minimized technical noise and ensured that subsequent analyses reflected biologically meaningful transcriptional patterns. The average total gene count per sample was 359,060 for control legs and 369,307 for loaded samples.

### Pathway enrichment analysis

Pathway enrichment analysis was performed using ShinyGO 0.85 [17] based on the lists of DEGs obtained from comparisons between the different AOIs. DEGs were mapped to curated gene sets from the Biological Process pathway database for *Mus Musculus* assembly with a false discovery rate of 0.05. Significant pathways were visualized using ShinyGO’s interactive tree and network plots to identify functional clusters and pathways.

### In vivo microCT

The right tibiae of the vehicle-treated, taurine-treated, *Slc13a5^cKO^*, and *Slc13a5^fl/fl^* mice were pre-scanned 4 days before first day of mechanical loading and post-scanned on day 18 using Scanco vivaCT40 (10.5 μm/voxel, 70 kVp, 114 μA, 300 ms integration) (Scanco Medical, Switzerland). For each tibia, a 2.1 mm region centered 5 mm proximally to the distal tibia-fibula junction was scanned while the mouse was under anesthesia with 1-3% isoflurane.

### In vivo scan registration and analysis

Pre and Post scans were registered in Dragonfly (Comet Technologies Canada Inc) using rigid registration, linear interpolation, and mutual information method. Registration results were exported as binary files and processed in MATLAB and BoneJ [18] to calculate 3D histomorphometry-like parameters. Mineralizing/eroding surface (MS, ES) were defined as the percentage of formation/resorption surface over the total periosteal or endosteal surface. Mineral apposition rate (MAR) and mineral resorption rate (MRR) were defined as the average thickness for formed/resorbed bone per day. Bone formation rate (BFR _BV_) and bone resorption rate (BRR _BV_) were calculated by dividing the volume of formed/resorbed bone by the total bone volume and number of days. All parameter calculations were based on previously described methodology [19].

Radial quantification was performed in MATLAB. First 3D image stacks from all samples were registered, and a common center was selected. Periosteal surface was discretized in 36 bins [20]. Areas of periosteal bone formation and resorption were quantified between each bin, slice by slice. The average measure per radial bin along the stack was calculated. Radial data was also grouped and analyzed based on anatomical locations, such as the posterior-lateral region (in compression), anterior-lateral region (in tension), and the medial and lateral regions (neutral axis).

### Ex vivo microCT

Prior to scanning, bones were rehydrated sequentially in 50% and 20% ethanol, followed by incubation in 1× PBS. Samples were embedded in 2% agarose to minimize movement artifacts and scanned using micro-computed tomography (µCT 50, Scanco Medical AG, Switzerland) at a voxel size of 10 µm/voxel, 70 kVp, 57 µA, and 700 ms integration time. MicroCT scans were analyzed using Scanco evaluation software (Scanco Medical AG, Switzerland). Cortical bone parameters, including cortical area (Ct.Ar), cortical thickness (Ct.Th), minimum moment of inertia (Imin), maximum moment of inertia (Imax), marrow area (Ma.Ar), total cross-sectional area (Tt.Ar), and cortical bone fraction (Ct.Ar/Tt.Ar), were quantified as previously described [21]. The cortical region of interest (ROI) was defined as a 5-mm segment located proximal to the distal tibia–fibula junction. For trabecular bone analysis, the ROI was defined as a 1-mm segment located distal to the proximal tibial growth plate. Trabecular parameters, including trabecular thickness (Tb.Th), trabecular number (Tb.N), trabecular separation (Tb.Sp), and bone volume fraction (BV/TV), were calculated using standard Scanco algorithms.

### *In vitro* Sclerostin treatment of primary osteoblast culture and quantitative RT-PCR

Primary calvarial osteoblasts (pCOBs) were isolated from postnatal day 1.5 pups as previously described [12], using sequential collagenase type I digestions, and expanded in α-MEM (Corning) supplemented with 10% FBS and 1% Penicillin-Streptomycin. After removal of non-adherent cells, cultures were maintained in growth medium, and experiments were performed using passage 1 or 2 cells. For osteogenic differentiation, cells were cultured in α-MEM supplemented with β-glycerophosphate (10 mM) and ascorbic acid (50 µg/mL) for 3 weeks prior to analysis. For sclerostin treatment, differentiated pCOBs were serum-reduced to 0.1% FBS and treated with 100ng/ml of recombinant mouse sclerostin (R&D systems) or vehicle control (PBS) for 24 h.

Total RNA was isolated from cultured cells following lysis in RLT buffer (Qiagen) using the RNeasy Micro Kit (Qiagen) in accordance with the manufacturer’s guidelines. For tissue samples, homogenization was performed in TRIzol reagent, followed by phase separation with chloroform and purification using the RNeasy Mini Kit (Qiagen). Reverse transcription was carried out using 1 µg of RNA and the iScript cDNA Synthesis Kit (Bio-Rad). Quantitative PCR analysis was conducted using iQ SYBR Green Supermix (Bio-Rad) and *Slc13a5*-specific primers were designed from the Harvard PrimerBank (ID: 281306805c1, sequence: F: ATGGATTCGGCGAAGACTTGT, R: AGGTATCAGAATGA CGAGTGGA). Amplification and detection were performed on a Bio-Rad real-time PCR system under standard cycling conditions. Transcript abundance was normalized to 18S rRNA, and relative expression levels were calculated using the comparative Ct (ΔΔCt) method.

### Statistics

Differential gene expression analysis of the spatial transcriptomic data was performed using a linear mixed-effects model (LMM), *p*-values <0.05 were considered statistically significant. Analysis was repeated for a set of pre-determined comparisons including: <u>Level1</u>: [Control vs Loaded]. <u>Level2</u>: [Control Bone vs Loaded Bone], [Control Periosteum vs Loaded Periosteum], [Control Periosteum vs Control Bone], [Loaded Periosteum vs Loaded Bone]. <u>Level3</u>: [Control Compression vs Loaded Compression], [Control Tension vs Loaded Tension], [Control Compression Periosteum vs Loaded Compression Periosteum], [Control Compression Bone vs Loaded Compression Bone], [Control Tension Periosteum vs Loaded Tension Periosteum], [Control Tension Bone vs Loaded Tension Bone], [Loaded Compression vs Loaded Tension], [Loaded Compression Periosteum vs Loaded Tension Periosteum], [Loaded Compression Bone vs Loaded Tension Bone] [Control Compression vs Control Tension], [Control Compression Periosteum vs Control Tension Periosteum], [Control Compression Bone vs Control Tension Bone].

The normality of the *Ex-vivo* microCT data was tested and parametric 2-way ANOVA was used to determine significant effects of mechanical loading and genotype on the cortical and trabecular bone parameters (p=0.05).

*In vivo* registration results were analyzed using two-sample unpaired t-test (Ps.Ms, Ps.Es, Ps.MAR, Ps.MRR, Es.Ms, Es.Es, Es.MAR, Es.MRR) to test the effect of genotype or treatment on the bone formation and resorption in the loaded legs. For radial quantification, mean formation and resorption area were computed within 36 angular bins for each specimen. Assumptions of sphericity and normality were not met with these data. Therefore, to test genotype effects along the radial profile while accounting for repeated measurements within subjects, generalized linear mixed-effects models (GLME) with a Gamma distribution and log link were used. Significance of fixed effects was assessed using Wald tests. To localize genotype differences, post-hoc analyses were performed independently for each bin using Gamma generalized linear models, followed by Benjamini–Hochberg false discovery rate (FDR) correction across bins (α = 0.05). For region-based analysis, bins were grouped into predefined anatomical regions (tension, neutral axis, and compression), and mean values per region were analyzed using Gamma generalized linear models with FDR correction across regions. All analyses were performed in MATLAB (Statistics and Machine Learning Toolbox). Two-sided tests were used, and statistical significance was defined as FDR-corrected p<0.05.

## Results

### A three-part analysis strategy facilitates region-specific evaluation of load-evoked changes in gene expression

The uniaxial compression tibia-loading model was used to induce changes in gene expression in the loaded right leg, while using the left leg as an internal control. Daily loading consisted of 60 compressive cycles at 4 Hz, with a target strain of -2200 µε in the posterior-lateral side (Fig.1A). This loading regimen has been previously shown to induce bone formation and alter gene expression in the mouse tibia [2], [5]. AOIs were manually defined in GeoMx software based on the mechanical stimulus distribution along the tibial midshaft under uniaxial compression (Fig.1B). Due to tibial geometry, this loading model produces a posterior-lateral region in compression and an anterior-medial region in tension. Accordingly, AOIs were defined to capture the compressive periosteum, compressive cortical bone, tensile periosteum, and tensile cortical bone (Fig.1C). We then analyzed the data at three hierarchical levels (Fig.1D). Level 1 compared gene expression between loaded and control legs after merging all AOIs within each leg. Level 2 first assessed differences in gene expression between periosteum and bone, and then evaluated loaded-versus-control differences separately within each tissue compartment. Level 3 examined the compressive and tensile regions independently, comparing gene expression between loaded and control legs within each mechanically defined AOI.

### Level 1 spatial analysis of merged AOIs identifies loading-induced gene changes that mirror bulk RNAseq

To validate regional spatial transcriptomics for detection of load induced changes in gene expression, we first merged all AOIs in the control limb and compared gene expression to the merged AOIs of the loaded limb. This simulates bulk RNAseq of bone after centrifugation to remove the marrow [5], [6], [23]. This identified 192 DEGs that were upregulated with loading and 108 that were downregulated (p<0.05; Log_2_FC >|0.5|). Pathway analysis of upregulated DEGs identified significant enrichment in ossification, bone development, bone mineralization, extracellular matrix organization, and vesicle transport with loading (Fig.2B). Consideration of GO terms related to bone (Supplemental Table 1) isolated 31 DEGs including upregulation of genes commonly associated with load-induced bone formation: *Col1a1*, *Bglap*, *Gja1*, *Sp7*, and *Alpl* (Fig.2A). Spatial transcriptomics datasets also showed downregulation of genes associated with bone remodeling and bone resorption a such as *Lgr4*, *Tgfbr3,* and *Notch2.* Other common markers of bone resorption such as *Mmp9*, *Ctsk*, *Acp5*, and *Tnfsf11,* trended down with loading but did not reach significance. The complete list is available in Supplemental Table 2.

**Figure 2:**
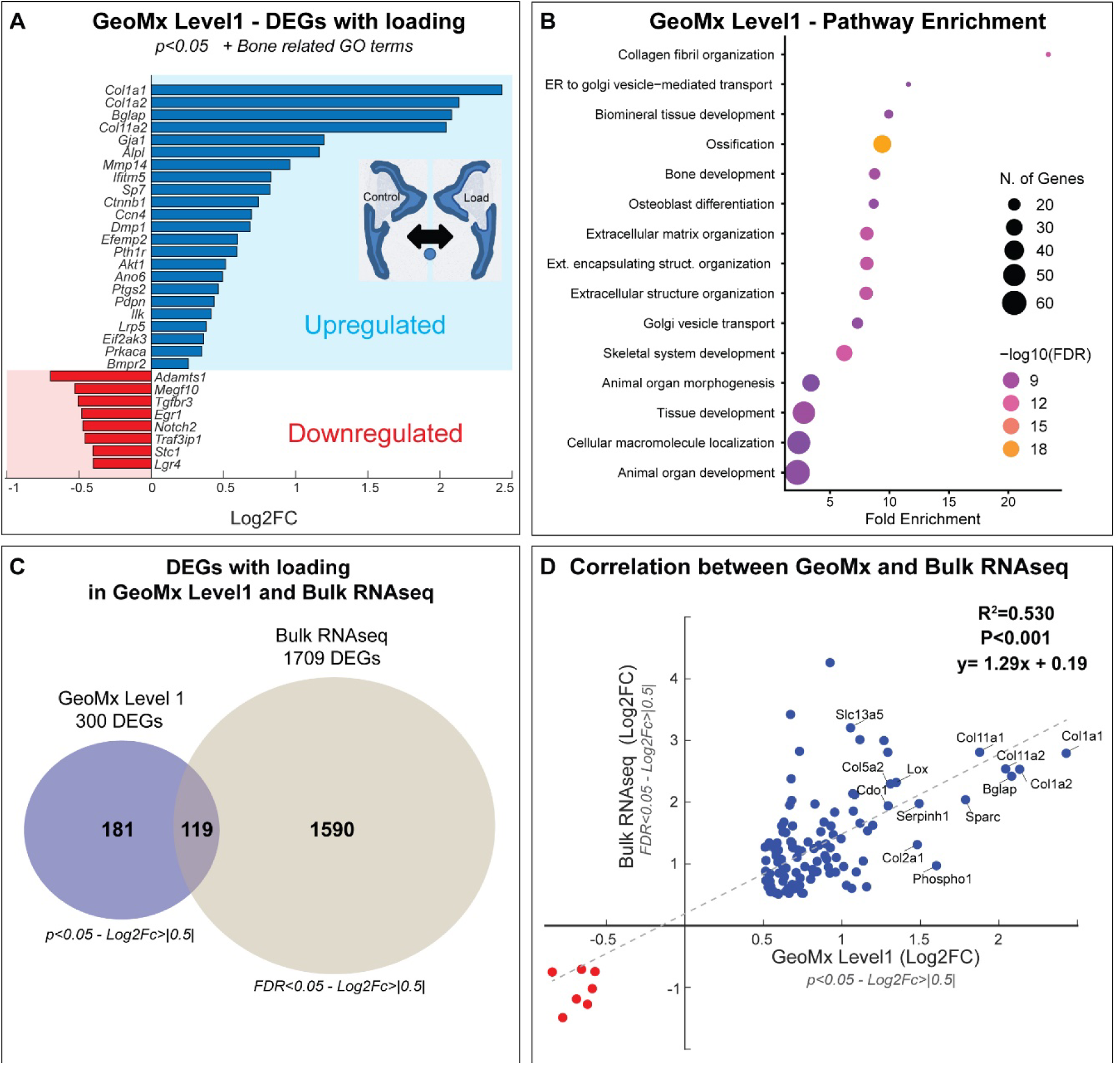
DEGs in the loaded samples compared to controls. All four periosteum and cortical bone AOIs were combined to mimic a bulk RNAseq analysis in marrow-depleted samples. A) Upregulated and downregulated genes with loading, filtered using Bone-related GO terms (Supplemental table 1) (p<0.05, Log2FC > |0.5|). Change in gene expression is reported as Log2FC. A positive Log2FC suggests an upregulation of the genes in the loaded legs. A negative Log2FC suggests a downregulation of the gene after loading. B) Pathway Enrichment analysis of enriched genes loading (ShinyGO 0.85). C) Comparison of the number of DEGs with loading between the GeoMx Level 1 analysis and publically available Bulk RNA sequencing data set from Chermside-Scabbo et al. [5]. D) Correlation of the Log2FC between shared DEGs between the GeoMx and bulk RNA sequencing data sets in (C).

Next, these findings were aligned with bulk RNAseq from an identical uniaxial tibia compression model and time course [5]. GeoMx and bulk RNAseq shared a total of 119 DEGs with loading (p<0.05; Log_2_FC>|0.5|; Fig.2C) among which, 112 genes were upregulated, and 7 were downregulated (Fig.2D). Linear regression identified a significant correlation in fold change of shared, loading-responsive genes between the two techniques (R^2^=0.530, p<0.001; Fig.2D). This included well-established osteogenic genes such as *Bglap*, *Col1a1*, *Col1a2*, *Gja1*, *Alpl*, and *Phospho1* (Fig.2D) in addition to less commonly reported genes such as *Col11*, *Serpinh1*, *Slc13a5,* and *Cdo1*.

### Level 2 spatial analysis shows robust resolution of gene signatures from periosteal and bone AOIs

The initial Level 2 spatial analysis examined genes expressed in bone versus periosteal AOIs from loaded and control legs. We hypothesized that genes for osteocytes would be enriched in bone while genes and pathways related to osteoblasts, fibroblasts, and vascular cells would be enriched in periosteum. Overall, in the control tibia, there were 1198 DEGs, 696 enriched in bone (including *Mepe*, *Wnt1*, and *Cd44*) and 502 enriched in periosteum (including *Postn*, *Col1a1*, and *Bglap*). Loading amplified this differential to 2034 DEGs (1150 in bone and 884 in periosteum); p<0.05; Log_2_FC >|0.5|) (Supplemental Table 3). Of these, periostin (*Postn,* +2.60 Log2FC) was the second most enriched gene detected in the periosteum behind collagen type III alpha 1 chain (*Col3a1*, +2.95 Log2FC). Markers of osteoblasts (*Bglap*, *Alpl*), extracellular matrix (*Col1a1*, *Col1a2*, *Dcn*, *Sparc*), osteoclasts (*Ctsk*), blood vessels (*Ptn*), and skeletal stem cells (*Prrx1*, *Ddr2*, *Itm2a*) were also enriched in periosteal AOIs (Fig.3A). Pathway analysis of periosteum-enriched DEGs showed enrichment for extracellular matrix organization, ossification, vascular development, and cell motility (Fig.3B).

**Figure 3:**
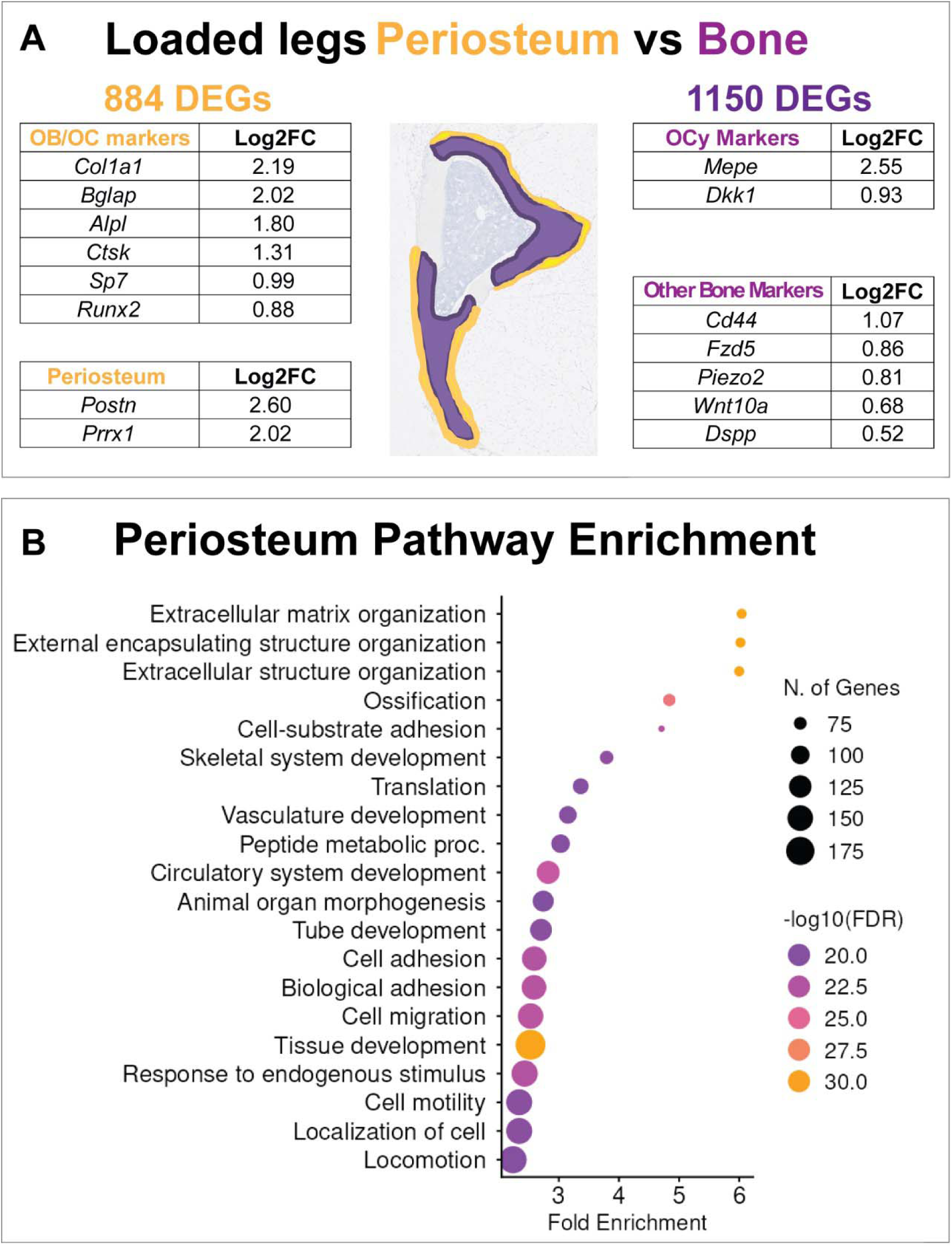
Level 2 analysis of periosteal vs bone AOIs. A) Differential expressed genes (DEGs) in the periosteum versus cortical bone of loaded legs. Common markers of osteoblasts, osteoclasts, periosteum, and osteocytes are reported with Log2FC. B) Pathway enrichment of DEGs higher in periosteum (ShinyGO 0.85).

Among top genes enriched in bone was matrix extracellular phosphoglycoprotein (*Mepe, +2.36 Log2FC*), a protein known to be highly enriched in mature osteocytes [24], and also Dickkopf-1 (*Dkk1*, Log2FC = 0.93) known as an inhibitor of the Wnt signaling pathaway (Fig.3A). Other markers such as transmembrane glycoprotein *Cd44*, Wnt-related genes *Wnt10a*, *Fzd5*, and mechanosensor *Piezo2* were also significantly enriched in bone AOIs (Fig.3A). Overall, GeoMx allowed for the detection of periosteal-and bone-specific gene profiles, with enhanced sensitivity in the loaded tibia.

### DEGs from bone and periosteal AOIs display some contamination with hematopoietic and muscle genes

One potential concern about the GeoMx platform is that it relies on laser-induced release of probe tags. Though AOIs were drawn within the boundaries of the cortical bone and periosteum, we hypothesized that there may be some non-specific gene contamination from neighboring bone marrow, blood vessels, and muscle, respectively. To check this, we first compared our dataset to the GO term hemopoiesis (GO:0030097) containing 1155 genes related to bone marrow and blood cells. Genes included in bone-related GO terms were subtracted (Supplemental table 1), leaving 129 “hemopoiesis” genes overlapping with our list of DEGs in bone vs periosteum (6.3% of total DEGs). Though many of these may be valid and represent immune populations within periosteum and cortical canals, genes enriched in the bone AOI also included hemoglobin subunit beta-2 (*Hbb-b2, 2.88 Log2FC)*, which is expressed mainly by erythrocytes, and *Igkc,* an immunoglobulin primarily expressed in B cells in the bone marrow (3.28 Log2FC). Muscle-related contaminants were similarly considered by identifying DEGs annotated under the GO terms “muscle system process” (GO:0003012) and “skeletal muscle tissue development” (GO:0007519). This included 652 genes, of which 79 overlapped with our list of DEGs (3.9% of total DEGs). Among these, 55/79 genes were enriched in the periosteum. This enrichment likely reflects contamination at the muscle-periosteum interface, where close physical proximity increases the risk of signal spillover. In some cases, expression of traditionally muscle-related genes including *Sox11*, *Hoxd9*, *Flt1*, *Map2k6*, and *Ddit3*, may be biologically plausable as these genes are also expressed in bone-related cell types. Despite this potential limitation, overall, >90% of DEGs matched the anticipated transcriptional signatures of the bone and perisoteum.

### Level 2 spatial analysis identifies unique loading-induced gene changes in bone and periosteal AOIs

The next Level 2 spatial analysis examined gene changes with loading in bone AOIs vs changes evoked specifically in periosteum. This identified 224 bone DEGs and 809 periosteal DEGs in response to mechanical load, with 41 gene changes shared between both regions including *Col1a1*, *Col1a2*, *Bglap, Phospho1, and Slc13a5* (Fig.4A,B; full lists in Supplemental Table 4). Pathway analysis indicated an upregulation of genes associated with bone mineralization within the cortical bone AOI, whereas genes linked to vesicle transport, osteoblast differentiation, and bone development were more prominently regulated in the periosteum (Supplemental Fig.1). This spatial segmentation of bone and periosteal tissues mirrors a prior study where laser capture microdissection was used to isolate periosteum and bone samples (referred to as Laser capture Mix) prior to RNA isolation and RNA sequencing [8]. Whereas laser capture required pooling multiple dissected fragments to yield adequate RNA, the GeoMx workflow achieved higher sensitivity and detected numerous additional DEGs from a single tissue section. Significant correlation in loading-induced gene changes was found between the laser capture and GeoMx periosteum data sets (Supplemental Fig.2).

**Figure 4:**
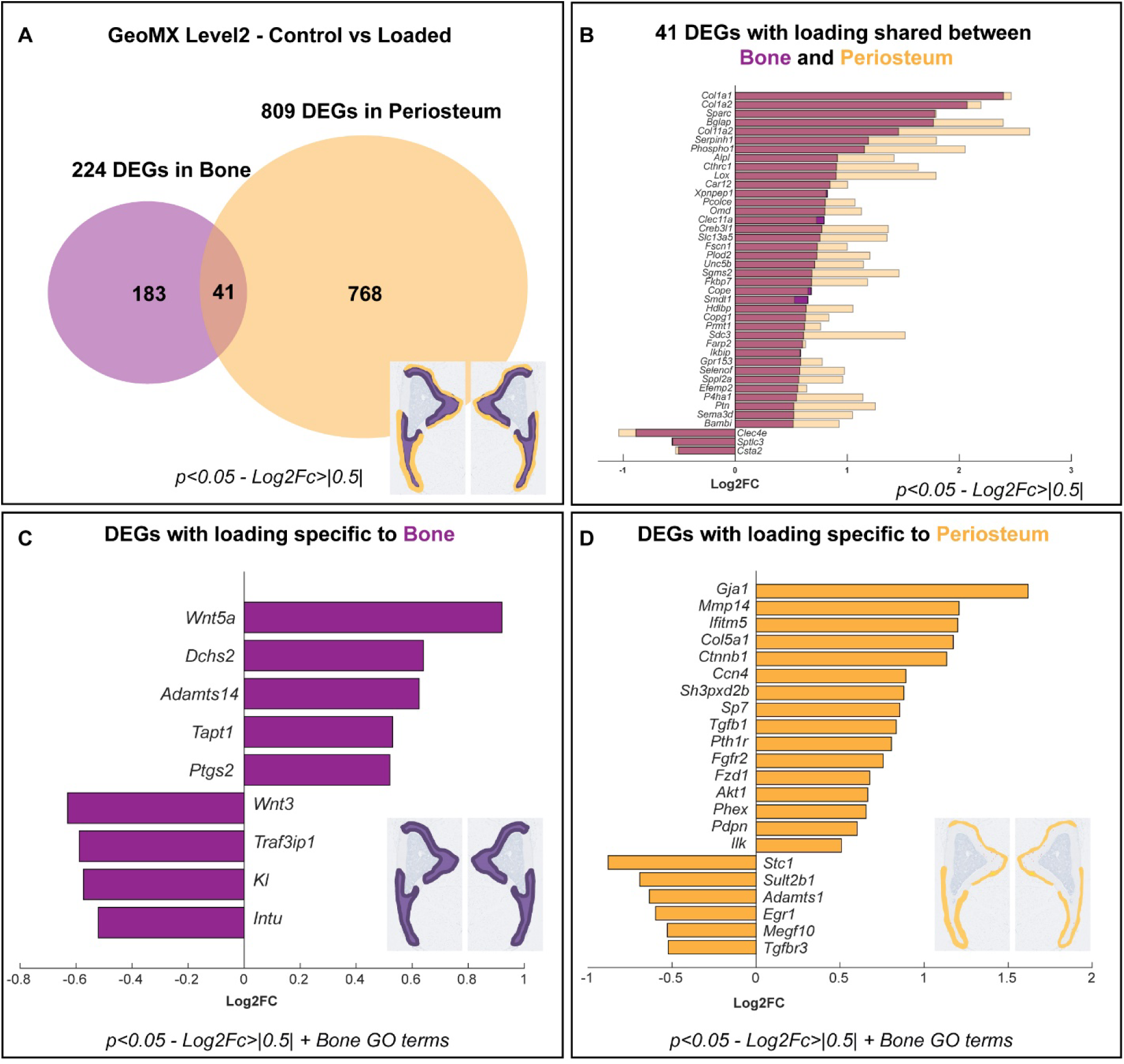
Sub-analysis of DEGs with loading in the periosteum and bone AOIs. Periosteum and cortical bone area were analyzed separately. A) DEGs with loading in the cortical bone, periosteum, and shared between the two AOIs. B) List of 41 DEGs with loading in both the periosteum (Orange) and the bone (Purple). Change in gene expression is reported as Log2FC. C) Bone-related DEGs with loading specific to the Bone AOI. D) Bone-related DEGs with loading specific to Periosteum AOI. A positive Log2FC suggests an upregulation of the genes in the loaded legs compared to control.

Specific to the bone AOI, *Ptgs2,* encoding for COX-2, was upregulated in agreement with a previous RNAseq data set [5]. *Ptgs2* is known to play a critical role in bone remodeling and in bone mechanotransduction [25], [26]. *Tapt1*, essential for trafficking proteins to the ciliary membrane of osteocytes, was also upregulated with loading. Mutations in *Tapt1* cause severe skeletal pathologies such as osteochondrodysplasia [27], highlighting its critical role in skeletal development and mechanotransduction. Downregulation of *Intu* and *Traf3ip1*, both involved in ciliary structure and function, was also noted. The Wnt signaling pathway was also affected with upregulation of non-canonical *Wnt5a* [28] and downregulation of canonical *Wnt3* with loading. *KI*, also known as Kotho, was also downregulated. Prior studies using Dmp1-Cre-mediated Klotho deletion reported increased bone formation, whereas Klotho overexpression suppresses mineralization *in vitro*, suggesting that reduced Klotho expression may favor anabolic responses to mechanical stimuli [29].

Specific to the periosteal AOI, extracellular matrix components including *Col5a1* were upregulated, consistent with increased collagen fibril organization during matrix deposition. Markers of osteoblast commitment and maturation, such as *Sp7*, *Phex*, and *Ifitm5* were also elevated, indicating a shift toward bone-forming phenotypes. Mechanotransduction mediators, including *Ilk*, were increased, consistent with enhanced force sensing at the bone surface. Genes related to several signaling pathways known to drive periosteal anabolism were also elevated including *Akt1*, *Ctnnb1*, *Fzd1*, and *Fgfr2*, pointing to coordinated engagement of Wnt and FGF signaling [30], [31]. Upregulation of *Pth1r* [32] and *Tgfb1* [33] also suggests heightened sensitivity to endocrine and paracrine cues that promote matrix synthesis and osteoprogenitor expansion. Increases in *Mmp14* [34] support localized matrix reorganization, while *Ccn4* likely facilitates osteoblast differentiation as a Wnt-induced matricellular cue [35]. Finally, upregulation of gap-junction gene *Gja1* is consistent with improved cell–cell communication during new bone formation and maturation toward osteocytes [31]. Conversely, we observed downregulation of periosteal genes including *Stc1*, which is involved in calcium and phosphate homeostasis [36], and *Adamts1*, which is involved in bone remodeling [37].

To summarize the Level 2 analysis, the GeoMx platform successfully distinguished bone from periosteum AOIs and enabled the discovery of load-induced gene expression changes specific to the periosteum and/or bone (full gene lists in Supplemental Table 3 & 4).

### Level 3 spatial analysis isolates unique load-induced gene adaptations at tensile vs compressive sites

During tibial loading, maximal bone formation occurs at the site of peak compression in the posterior-lateral region of the tibia [2]. By contrast, the site of peak tension in the anterior-medial region does not accrue as much bone. We hypothesized that these differences reflect distinct transcriptional responses to mechanical loading. To test this, we compared loading-induced gene expression changes (control vs loaded legs) in compressive and tensile regions using merged bone and periosteum AOIs. This analysis identified 580 load-induced DEGs on the compressive side and 189 on the tensile side (Fig.5A, Supplemental Table 5).

**Figure 5:**
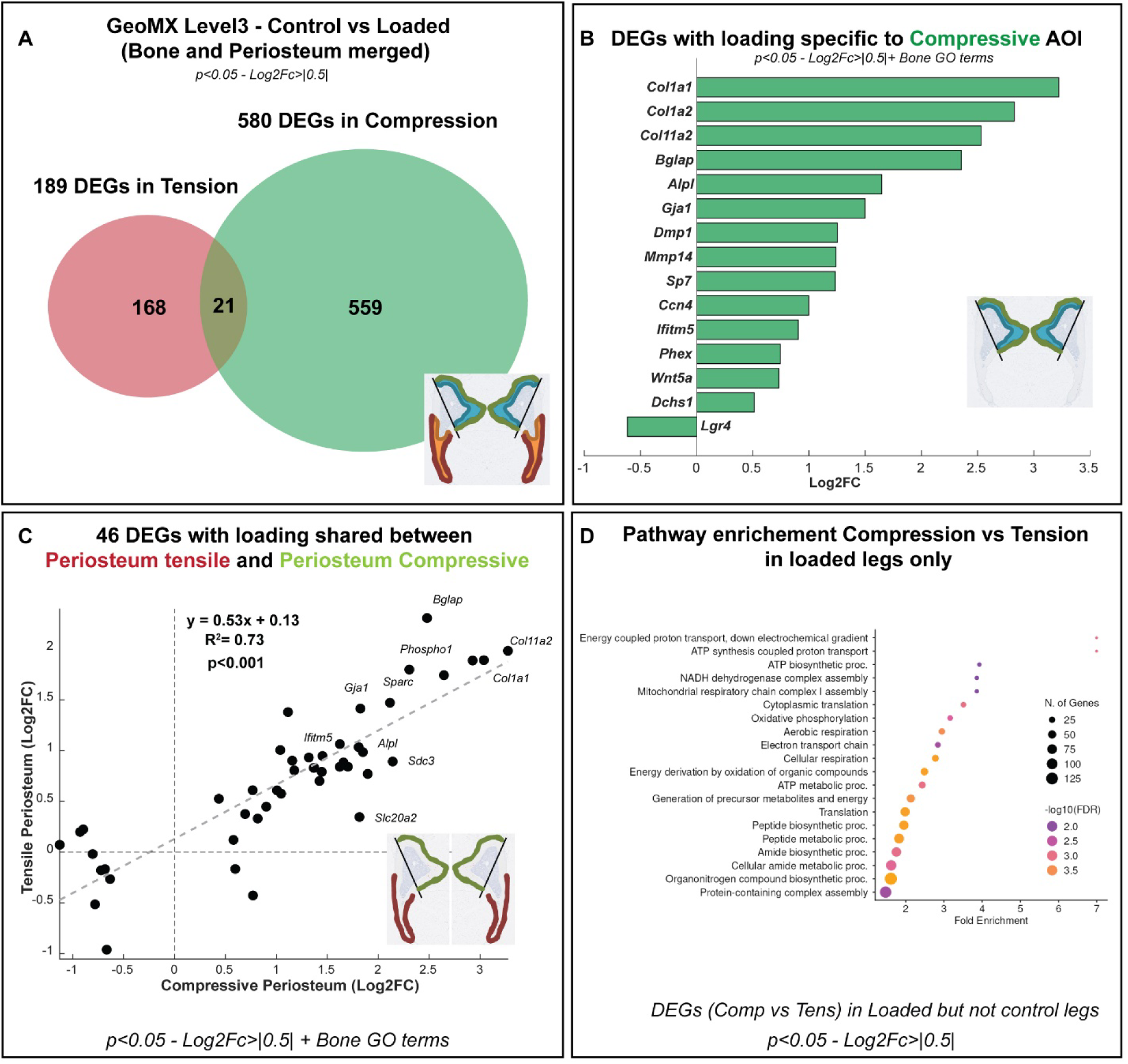
GeoMx Level 3 analysis comparing effect of loading on gene expression from the compressive and tensile region of the tibiae. A) DEGs with loading in the compressive and tensile AOIs (periosteum and bone merged). B) Regulation of Bone-related DEGs with loading in the compressive AOI. C) Correlation of DEGs shared between compressive and tensile periosteum. D) Pathway enrichments from genes found to be differentially expressed between compression and tension in the loaded leg but not in control leg.

Among these, 21 genes were regulated by loading in both regions, with highly correlated fold changes between compressive and tensile sites (R²=0.92, p<0.001; slope=0.73) (Supplemental Fig.3). Shared genes included upregulation of osteoblast-associated genes (*Bglap*, *Col2a1*, *Phospho1*, *Lox*) and downregulation of genes involved in mitochondrial and nucleotide metabolism (*Slc25a36*, *Gmpr*) and Wnt signaling (*Lgr4*). Tensile-specific DEGs (168/189) included upregulation of *Ptgs2* and downregulation of *Stc1* and *Camk2d*, but did not show significant pathway enrichment. In contrast, compressive-specific DEGs (559/580) included osteogenic and matrix-related genes (*Col1a1*, *Alpl*, *Gja1*, *Dmp1*, *Sp7*, *Phex*, *Mmp14*, *Ccn4*, *Wnt5a*, *Ifitm5*) and were enriched for pathways related to collagen fibril organization, matrix development, vesicle transport, and cellular localization (Fig.5B & Supplemental Fig.4).

Notably, 43% of compressive DEGs overlapped with periosteal DEGs identified in the Level 2 analysis, indicating that a substantial component of the compressive transcriptional response originates from the periosteum. Focusing on periosteum alone, we identified 46 DEGs induced by loading specifically in the compressive periosteum (Fig.5C). Comparison of fold changes for these genes between compressive and tensile periosteum revealed a strong correlation (R²=0.73, p<0.001, slope=0.53) with an approximately twofold greater magnitude of regulation on the compressive side, consistent with stronger local mechanical stimulation and enhanced mechanotransduction.

To further isolate intrinsic biological differences between compressive and tensile periosteum while minimizing potential muscle contamination, we compared DEGs between compression and tension in loaded legs after excluding genes that were also differentially expressed in control legs. This internally controlled analysis revealed strong enrichment of mitochondrial and metabolic pathways in the compressive periosteum, including oxidative phosphorylation, ATP synthesis, and NADH dehydrogenase complex assembly (Fig.5D), even after filtering muscle-associated genes, consistent with recent bulk RNAseq data analysis [38]. In contrast, the tensile periosteum was enriched for genes associated with matrix remodeling and osteoclast-related processes (*Ctsk*, *Tnfsf11*, *Mmp13*, *Mmp9*), as well as signaling pathways (*Wnt3*, *Spp1*, *Shh*, *Hoxd4*), without comparable metabolic activation.

Collectively, these Level 3 analyses demonstrate that tibial loading elicits highly region-specific transcriptional responses aligned with local mechanical environments. Compressive regions, particularly within the periosteum, engage robust osteogenic and metabolically intensive gene programs, whereas tensile regions activate remodeling-and turnover-associated pathways. These spatially resolved transcriptional differences provide a molecular framework linking mechanical stimulus magnitude to differential periosteal adaptation and help explain preferential bone formation at sites of peak compression.

### GeoMx spatial transcriptomics identifies Slc13a5 as a candidate mechanoresponsive gene

Our GeoMx Level 1 dataset was compared to three publicly available data sets to identify consistent DEGs with loading [5], [6], [8]. Only 12 genes were differentially expressed with loading in all four data sets (Fig.6). The most highly upregulated gene was *Col1a2* (+2.13 Log2FC). Among the rest, several fell into pathways that are less commonly discussed in the context of mechanical loading but appear relevant to skeletal adaptation. *Smpd3*, a lipid-metabolizing enzyme expressed in bone and cartilage, is required for normal skeletal development. Early developmental conditional deletion in chondrocytes and osteoblasts causes skeletal deformities, whereas post-developmental deletion results in only mild defects [39]*. Cthrc1*, thought to be a downstream target of BMP2, influences bone mass, with knockout models exhibiting low bone mass and overexpression producing a high-bone-mass phenotype [40]. We also detected factors such as *Sema3d*, which might modulate osteoclast activity through TNF-α signaling [41]. Several of these genes or their associated pathways have been implicated in human GWAS for bone mineral density or skeletal traits, such as *Smpd3, Sema3d, Cdo1*, or *Knck1* underscoring their relevance to human bone biology.

**Figure 6:**
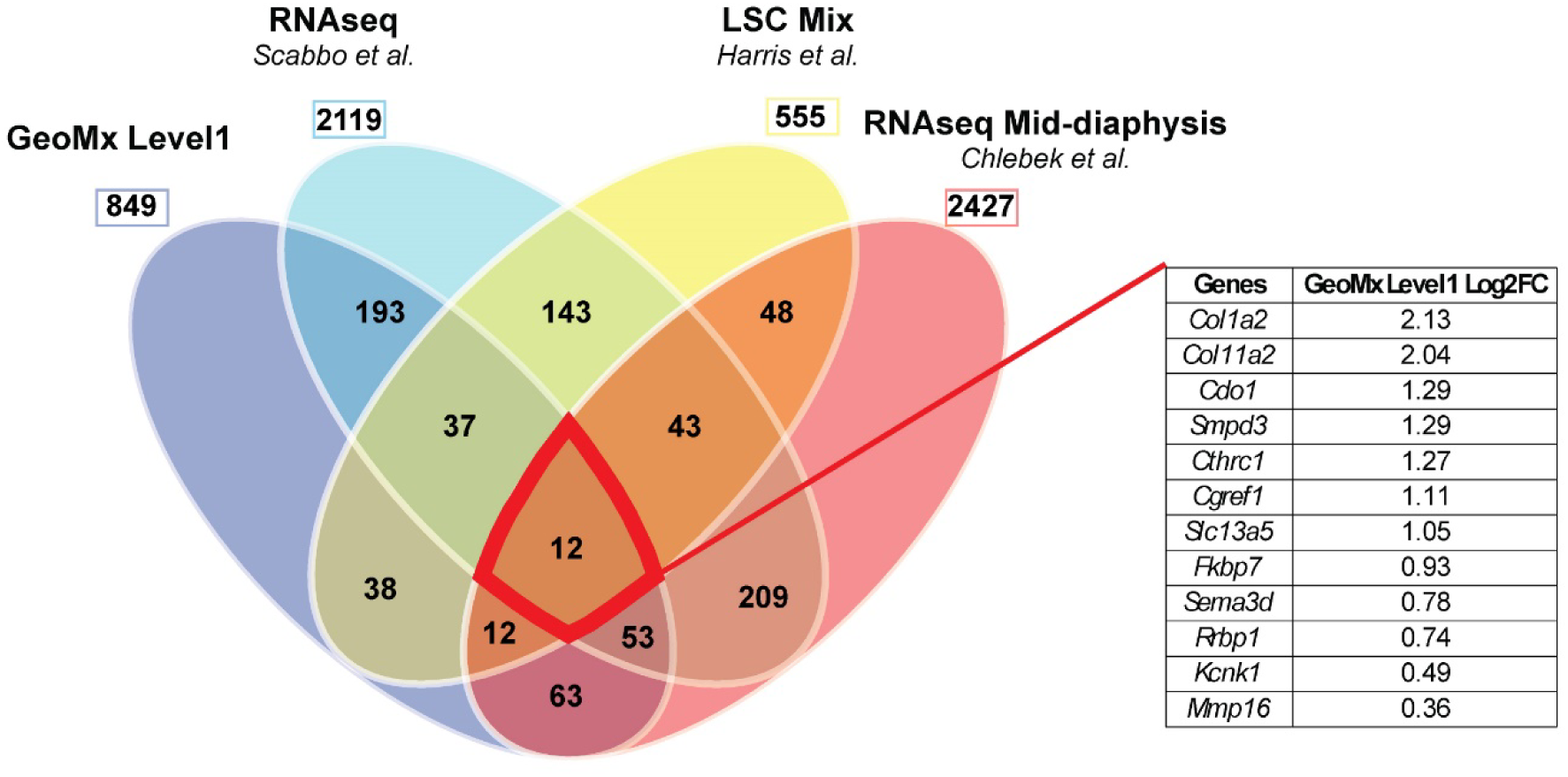
Venn Diagram comparing DEGs from four different studies investigating the effect of loading on bone gene expression. A) DEGs in our GeoMx Level1 study (p<0.05), compared to RNAseq (FDR<0.05) and laser capture (p<0.05) datasets. A total of 12 genes were differentially expressed with loading in all four studies. Gene names are listed in the table and Supplemental table 6.

Among the genes consistently regulated by loading, *Slc13a5* stood out as a candidate for further consideration given its function as a sodium-dependent citrate transporter and its reported involvement in bone mineralization [12]. Prior drug-target Mendelian randomization analyses indicated that genetic variants predicting reduced *Slc13a5* function were associated with decreased osteoporosis risk, based on both clinical diagnoses and self-reports in the UK Biobank cohort [42]. These findings suggest that *Slc13a5* inhibition may represent a potential strategy to mitigate age-related bone loss. However, the role of *Slc13a5* in bone mechanoadaptation remains unexplored. In our spatial transcriptomics data set, *Slc13a5* was enriched in the loaded legs (Level1, +1.05 Log2FC, *p*=0.02). Level 2 analysis showed that *Slc13a5* was significantly enriched in the loaded periosteum compared with control (+1.35 log2FC, *p* < 0.01). It was also enriched in loaded bone (+0.76 log2FC, *p* < 0.01). This pattern is consistent with its established expression in late osteoblasts and early osteocytes [12].. Level 3 analysis showed that *Slc13a5* was enriched in the compressive side with loading (Bone+Perio AOIs merged, +1.51 Log2FC, *p*<0.005). Thus, we selected *Slc13a5* for targeted functional investigation.

### Conditional deletion of osteoblastic Slc13a5 enhances bone formation and reduces resorption in regions of low mechanical stimulation

*Slc13a5* is a sodium-dependent citrate transporter expressed on the plasma membrane of various cell types, including osteoblasts [12]. Citrate plays a central role in cellular energy metabolism via the TCA cycle and glycolysis [43], and the majority of the body’s citrate is stored in bone [44]. Although global knockout of *Slc13a5* was reported to protect aged mice against adiposity and insulin resistance [11], global deletion of this gene can also lead to abnormal bone structure, reduced mineral density, and bone strength. However, recent studies showed that conditional deletion of *Slc13a5* in osteolineage cells with Ocn-Cre has differential effect in young growing mice and adult mice. Indeed, in adult mice, conditional osteoblast deletion reduces tissue mineral density but does not affect bone structural parameters [12], [42]. Mendelian randomization analysis of UK Biobank data indicated that variants predicted to reduce *Slc13a5* function are associated with decreased osteoporosis risk [42]. These suggest that inhibition of this citrate channel in osteoblasts could be beneficial for bone quality with aging.

To investigate whether *Slc13a5* in the late osteolineage is necessary for bone mechanoadaptation, we analyzed the response to loading in Ocn-Cre *Slc13a5* conditional knockout (*Slc13a5^cKO^*) mice compared to *Cre* negative *Slc13a5^fl/fl^* control littermates. Using traditional *ex vivo* microCT analysis of loaded versus nonloaded control tibias, we observed a significant effect of loading in both genotypes. Parameters including Ct.Th, Ct.Ar, Imin, and Tb.Th were all significantly increased with loading (Supplemental Fig.6). *Slc13a5^cKO^* mice exhibited lower bone mineral density (BMD) (Supplemental Fig.6) compared to Cre- littermates as previously reported [12], [42].

To investigate potential spatial effects of the conditional deletion, we used time-lapse *in vivo* microCT analysis to capture bone formation and resorption metrics in 3D, with subsequent radial quantification of loaded tibiae from both genotypes. This showed that both *Slc13a5^cKO^* and control mice responded to mechanical loading, with bone formation localized primarily at the site of peak compression (Fig.1B) in the posterior-lateral region of the tibia (Fig.7A). In loaded limbs of *Slc13a5^cKO^*mice, periosteal mineralizing surface was 16% greater compared to loaded limbs of control mice, while eroding surface was 52% less (Fig.7B). This difference in resorption was also reflected in the 3D rate parameter, BRR_(BV)_ (*Cre- Slc13a5^fl/fl^*: 0.03%/day; *Slc13a5^cKO^*: 0.016%/day, *p*=0.02). On the other hand, 3D formation rate, BFR_(BV)_ was found to be similar between genotypes (*Cre- Slc13a5^fl/fl^*: 0.29%/day; *Slc13a5^cKO^*: 0.29%/day). Similar observations were made for rates of mineral apposition (MAR) and resorption (MRR) (Fig.7B). These results indicate that the overall amounts of bone formation and resorption were comparable between genotypes, but a greater proportion of the bone surface was undergoing formation (based on MS/BS), and a smaller proportion was undergoing resorption (based on BRR_(BV)_), in *Slc13a5^cKO^* compared with controls. This suggests a spatial redistribution of remodeling activity.

**Figure 7:**
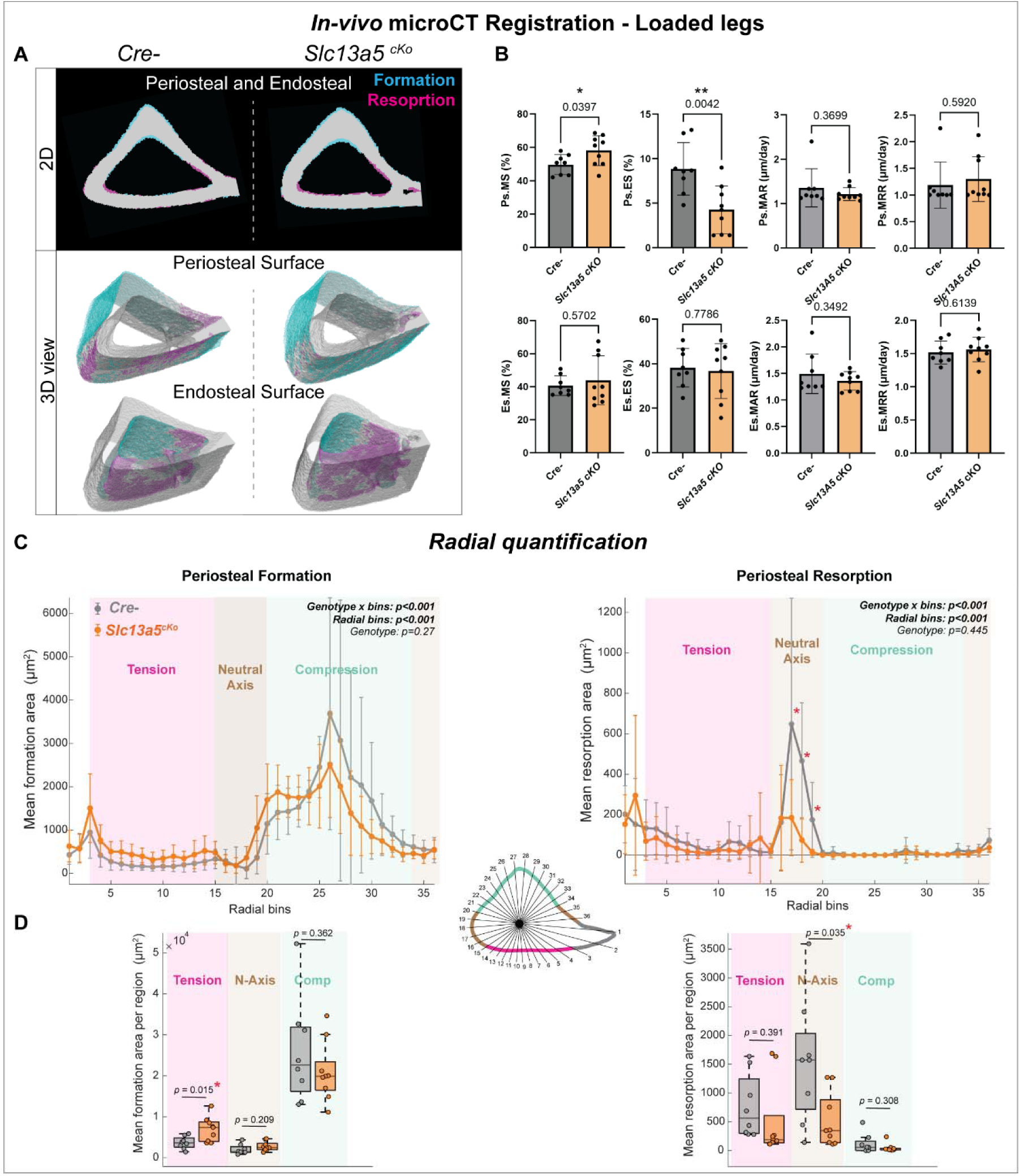
Effect of conditional deletion of osteoblastic *Slc13a5* on the bone response to loading. A) *In* vivo microCT registration and analysis. Pre and post scans were registered. Regions of formation (cyan) and resorption (magenta) are shown in 2D and 3D on the periosteal and endosteal surfaces, for both genotypes. Histomorphometry-like parameters were calculated based on the 3D registration. Radial analysis shows the amount of bone formation and resorption around the periosteal surface. (*Slc13a5 ^cKo^*: n=9, Cre-: n=8).

To investigate this, we performed a radial analysis. This approach identified a significant increase in bone formation along the anterior-medial region (under tension) of the *Slc13a5^cKO^*tibiae (Fig.7C,D), and reduced resorption along the neutral axis compared to *Slc13a5^fl/fl^*controls (Fig.7C,D). Together, these findings suggest that conditional knockout of *Slc13a5* facilitates bone mechanoadaptation in regions of the tibia experiencing low-magnitude mechanical stimulation.

Lastly, to explore a potential mechanistic link between bone cell mechanoregulation and *Slc13a5*, we examined the effect of the mechanoresponsive Wnt antagonist, sclerostin, on *Slc13a5* expression *in vitro*. Treatment of primary osteoblast cultures with recombinant sclerostin resulted in a significant decrease in *Slc13a5* expression (Fig.8), suggesting that osteocyte-derived signals can directly modulate *Slc13a5* levels in osteolineage cells. This finding complements our spatial transcriptomics results showing that *Slc13a5* is upregulated with mechanical loading, a condition known to reduce sclerostin production, and supports a model in which osteocytes regulate *Slc13a5* in a load-dependent manner.

**Figure 8:**
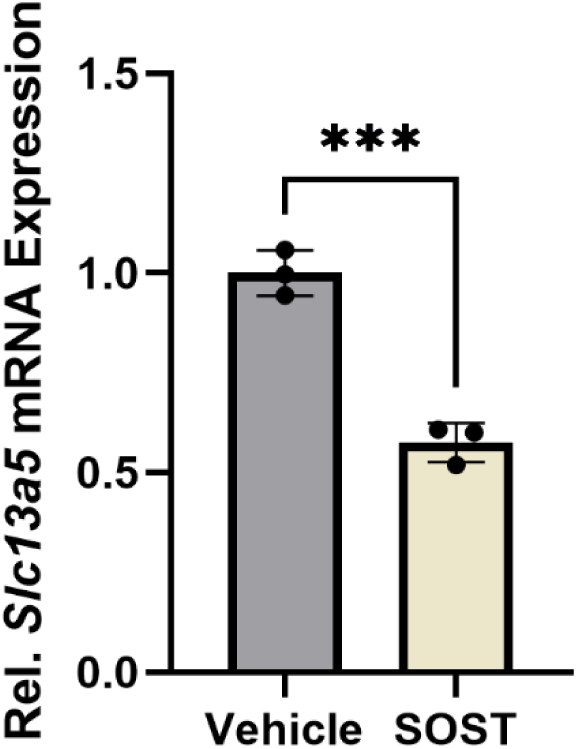
Relative expression of *Slc13a5* in primary osteoblasts culture treated with recombinant sclerostin (SOST) or PBS vehicle (Veh). Graph represents mean±SD; Student’s t-test versus control, ***p<0.001, (n=3).

## Discussion

In this study, we demonstrated the utility of the GeoMx spatial transcriptomics platform for investigating spatial regulation of gene expression following mechanical loading in bone. We examined differences between periosteum and cortical bone as well as spatially distinct transcriptional responses at compressive and tensile surfaces. The approach was validated through multi-level analyses, including comparisons with publicly available bulk RNA-sequencing datasets. The spatial resolution provided by GeoMx enabled detection of gene expression differences within and between loaded and contralateral limbs across defined bone subregions, including periosteum, cortical bone, and regions experiencing compression or tension.

This subregion-specific resolution was a key advantage of the GeoMx approach and revealed substantially more DEGs in Level 2 analyses than in Level 1 analyses, in which subregions were merged. Level 2 results showed strong concordance with publicly available laser capture microdissection datasets, further validating the spatial framework. Notably, many loading-responsive transcriptional changes are localized to the compressive surface of the tibia, predominantly within the periosteum, as revealed by the Level 3 analysis.

We detected genes likely reflecting partial signal contamination from adjacent tissues, including bone marrow and muscle (<10% of genes). Bone AOIs may capture transcripts from neighboring bone marrow or hematopoietic cells within transcortical vascular canals, whereas periosteal AOIs are more susceptible to inclusion of muscle-derived RNA. These observations highlight the complexity of spatial transcriptomic datasets, in which transcripts may be shared across cell populations, complicating the distinction between biological signals and partial tissue overlap. Gene-filtering strategies can therefore be applied depending on study objectives and, in this study, we were successful in identifying many relevant DEGs. Another limitation is that increasing subregion specificity (*e.g.*, comparing compressive versus tensile periosteum) reduces the size of the analyzed regions, which in turn lowers total gene counts and may decrease the sensitivity of the outputs.

Despite limitations, our spatial approach identified novel candidate regulators of mechanoadaptation, including *Slc13a5*. Functional analysis of *Slc13a5* revealed reduced load-induced bone resorption near the neutral axis and increased bone formation in tensile regions in osteoblastic conditional knockout mice compared to controls. The conditional deletion of *Slc13a5* predominantly affected areas experiencing low mechanical stimulation. This pattern contrasts with transcriptomic findings showing loading-induced upregulation of *Slc13a5* primarily in the compressive region, the site of maximal bone formation. This discrepancy suggests that *Slc13a5* may not act as a primary driver of osteogenesis at highly loaded sites, but rather as a metabolic modulator that constrains remodeling in mechanically permissive or low-magnitude environments. Upregulation of *Slc13a5* on the compressive surface may reflect increased metabolic demand during active bone formation [38], [43], whereas its deletion alters remodeling where mechanical cues are weaker. Alternatively, *Slc13a5* deficiency may modify baseline bone structure or cellular metabolism, increasing skeletal sensitivity to loading and enabling adaptation at lower stimulus magnitudes. Loss of *Slc13a5* would thus lower the mechanical threshold for adaptation, revealing effects in regions normally below the osteogenic activation threshold. Notably, *Slc13a5* is expressed in late osteoblasts and early osteocytes [12], which detect mechanical stimuli and coordinate osteoblast and osteoclast activity through paracrine signaling. Expression of *Slc13a5* in these cells may therefore influence the sensitivity or set point of mechanoadaptive responses across bone microenvironments. Osteocytes are known to express *Sost*, a key regulator of bone mechanoadaptation whose expression is reduced by mechanical loading [4], [9], [45]. In the present study, the inhibitory effect of sclerostin treatment on *Slc13a5* expression in osteoblast cultures supports the hypothesis that *Slc13a5* is regulated, at least in part, by osteocyte-derived signals.

Such a mechanism is particularly relevant in osteoporosis, where impaired mechanoadaptive capacity leads to low bone mass and increased fracture risk. Strategies that lower the mechanical threshold required to trigger bone formation could represent a promising therapeutic approach to enhance skeletal adaptation and prevent bone loss.

## Conclusions

In summary, this study establishes spatial transcriptomics as a robust and informative approach for investigating bone mechanoadaptation. By resolving transcriptional responses within the periosteum, cortical bone, and mechanically distinct regions of the tibia, we demonstrate that compressive loading elicits a uniquely strong, osteogenic, and metabolically demanding gene program, particularly within the periosteum. This spatially biased response provides a molecular framework for understanding preferential bone formation at sites of peak mechanical stimulus. Furthermore, our integrative analysis identifies *Slc13a5* as a novel metabolic regulator linked to mechanical loading, highlighting connections between energy metabolism and skeletal mechanobiology. Together, these findings expand current models of bone adaptation to mechanical forces and provide a valuable resource for identifying new therapeutic targets to enhance bone formation and preserve skeletal health.

## Data Availability

The full dataset generated in this study is available at [GSE324027].

## Supporting information

Supplemental Results

## Acknowledgments

We would like to thank David DeBruin (Saint Louis University), The Research Microscopy & Histology Core (Saint Louis University), and Thomas Andersen (University of Southern Denmark) for their input and support.

## Author Contributions

**Quentin A. Meslier:** Conceptualization - Formal analysis – Investigation – Visualization-Writing - Original Draft - Writing - Review & Editing. **Alec T. Beeve:** Methodology – Validation – Investigation. **Arushi Gupta:** Formal analysis. **Daniel Palomo:** Formal analysis. **Samia Saleem:** Formal analysis – Visualization. **Serena Eck:** Methodology – Resources. **Lisa Lawson:** Validation – Investigation. **John Shuster:** Validation – Investigation. **Michelle Brennan**: Data Curation – Methodology – Software. **Naomi Dirckx:** Methodology – Resources - Writing - Review & Editing - Funding acquisition. **Matthew J. Silva:** Conceptualization – Methodology – Resources - Writing - Review & Editing - Supervision - Funding acquisition. **Erica L. Scheller:** Conceptualization – Methodology – Visualization - Writing - Original Draft - Writing - Review & Editing – Supervision - Funding acquisition

---

**Supplemental Figure 1:**
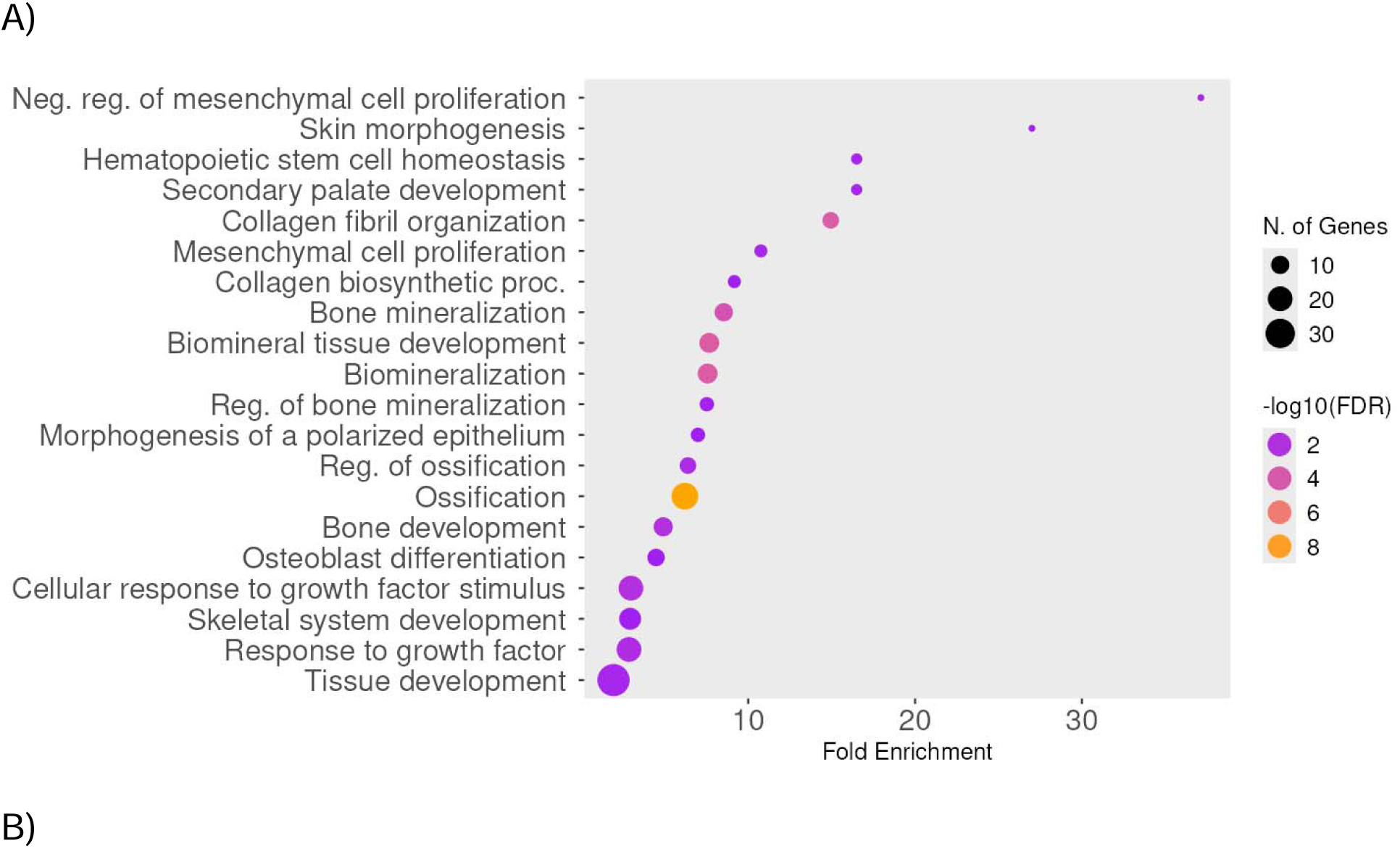
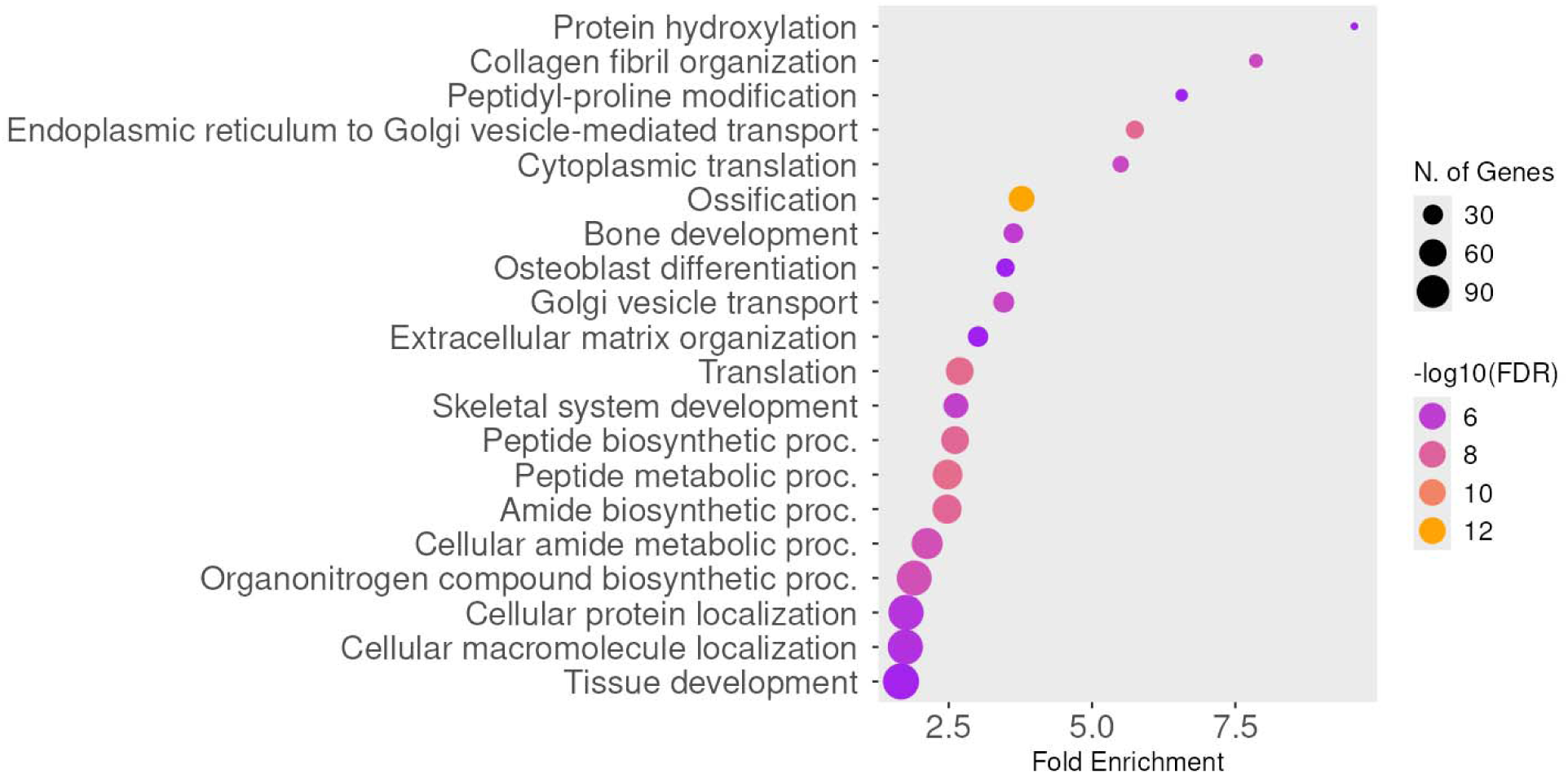
Pathway Enrichment analysis of GeoMx Level 2 analysis results that are enriched with loading. A) In the cortical bone (Control Bone vs Loaded Bone) B) In the periosteum (Control Perisoteum vs Loaded Perisoteum) (ShinyGO 0.85).

**Supplemental Figure 2:**
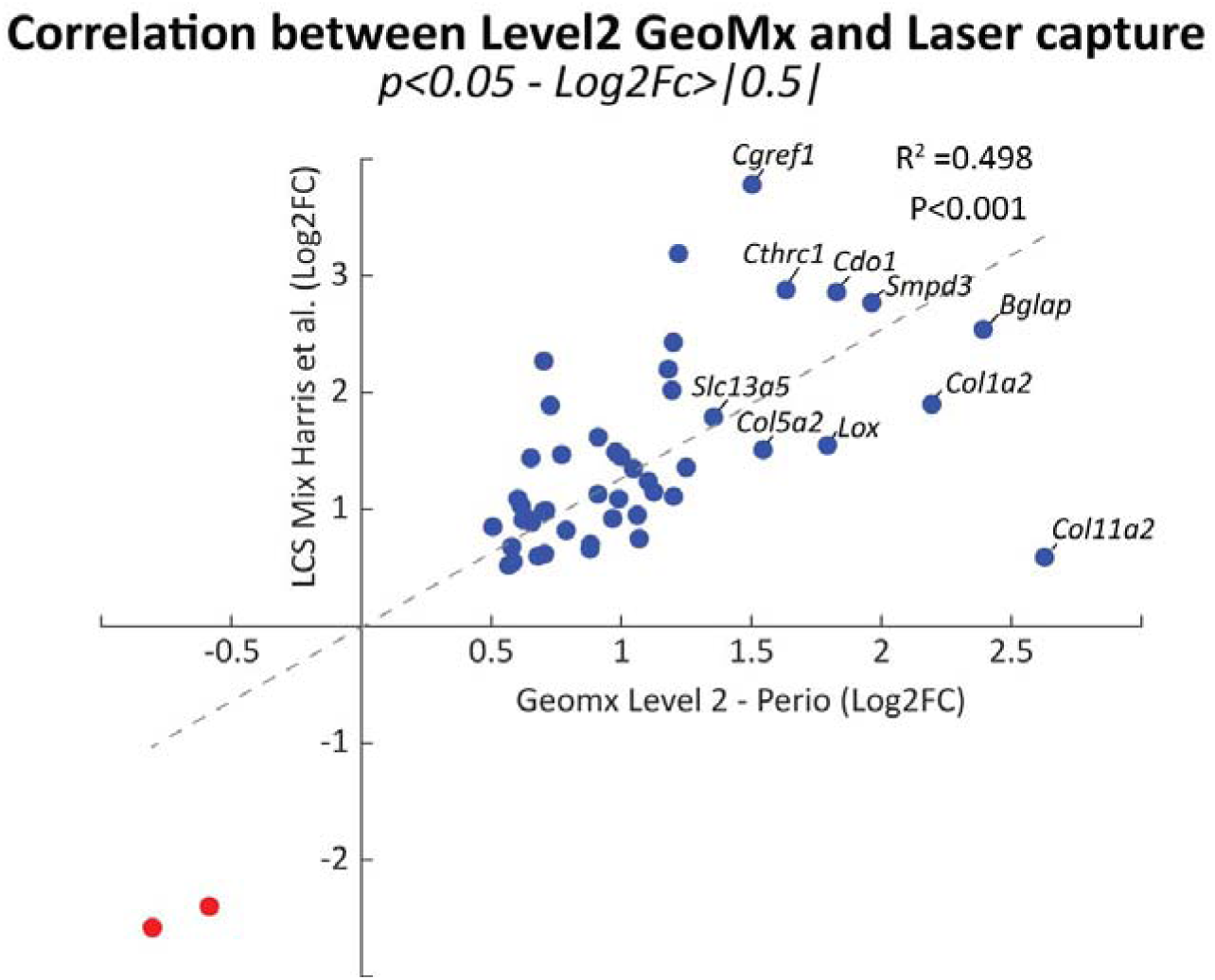
Correlation between GeoMx Level 2 data set results (Control Periosteum vs Loaded Periosteum) and microdissection laser capture data set (Harris et al.)

**Supplemental Figure 3:**
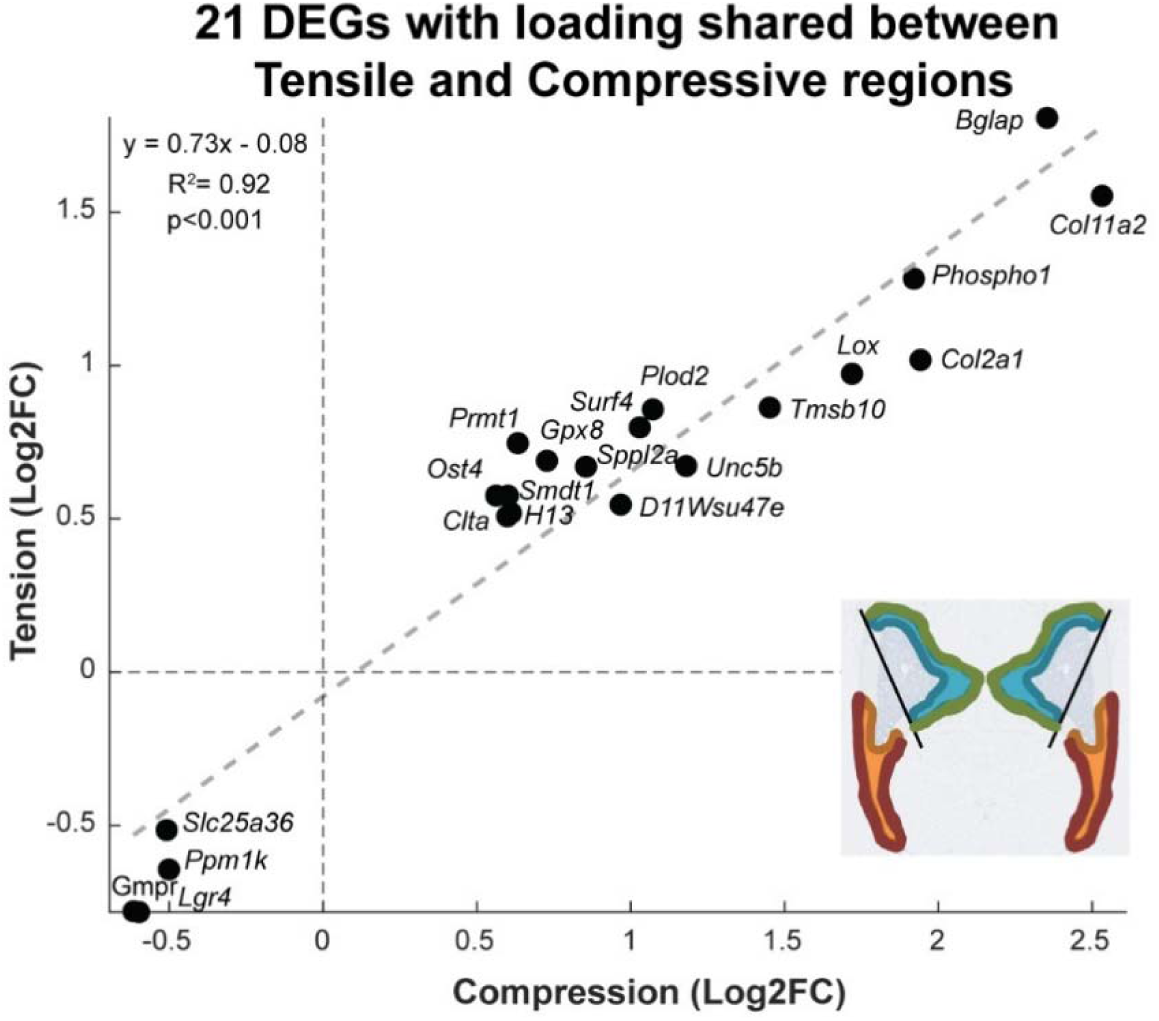
Correlation between GeoMx Level 3 genes enriched in loaded compressive regions (Perio/Bone merged) and genes enriched in loaded tensile regions (Perio/Bone merged).

**Supplemental Figure 4:**
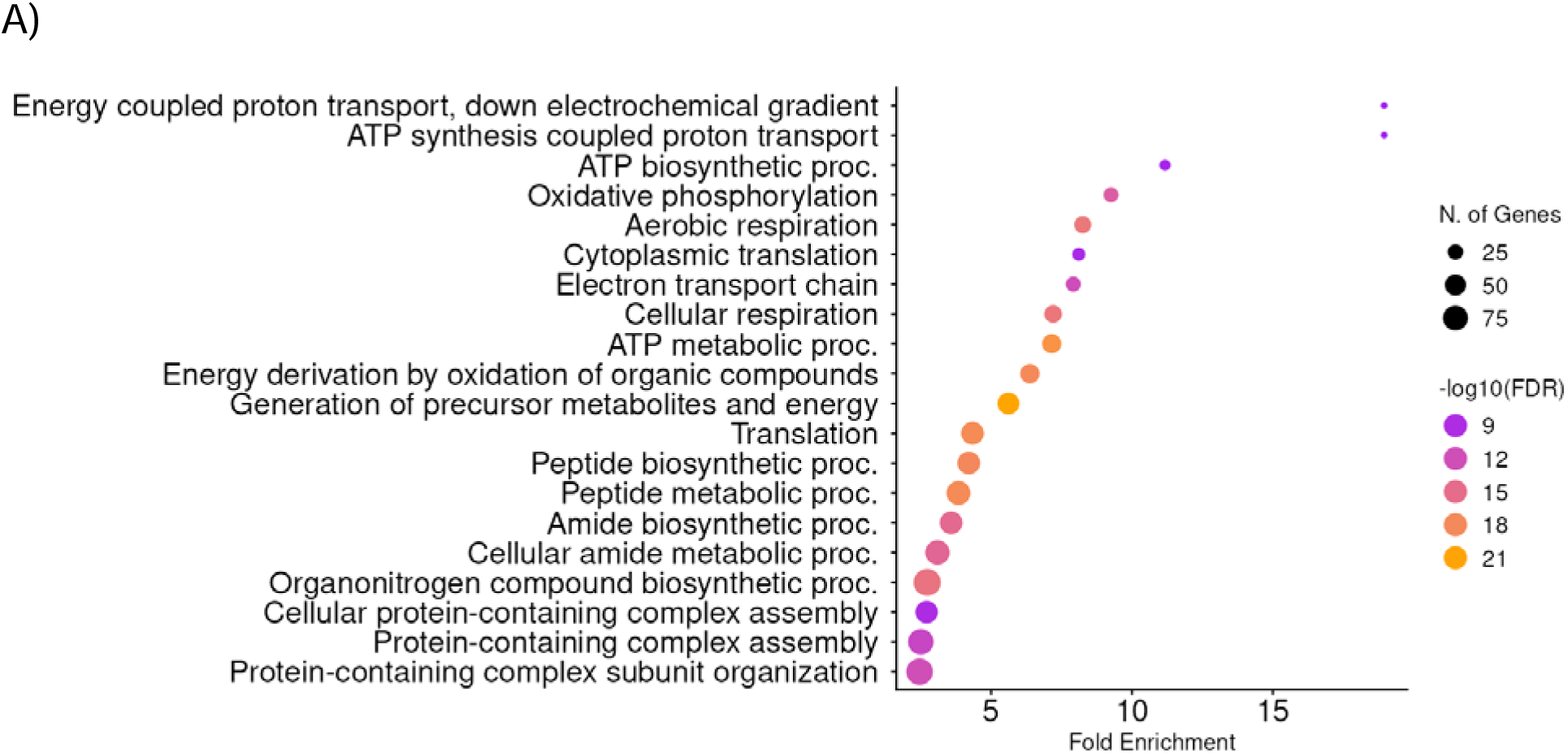
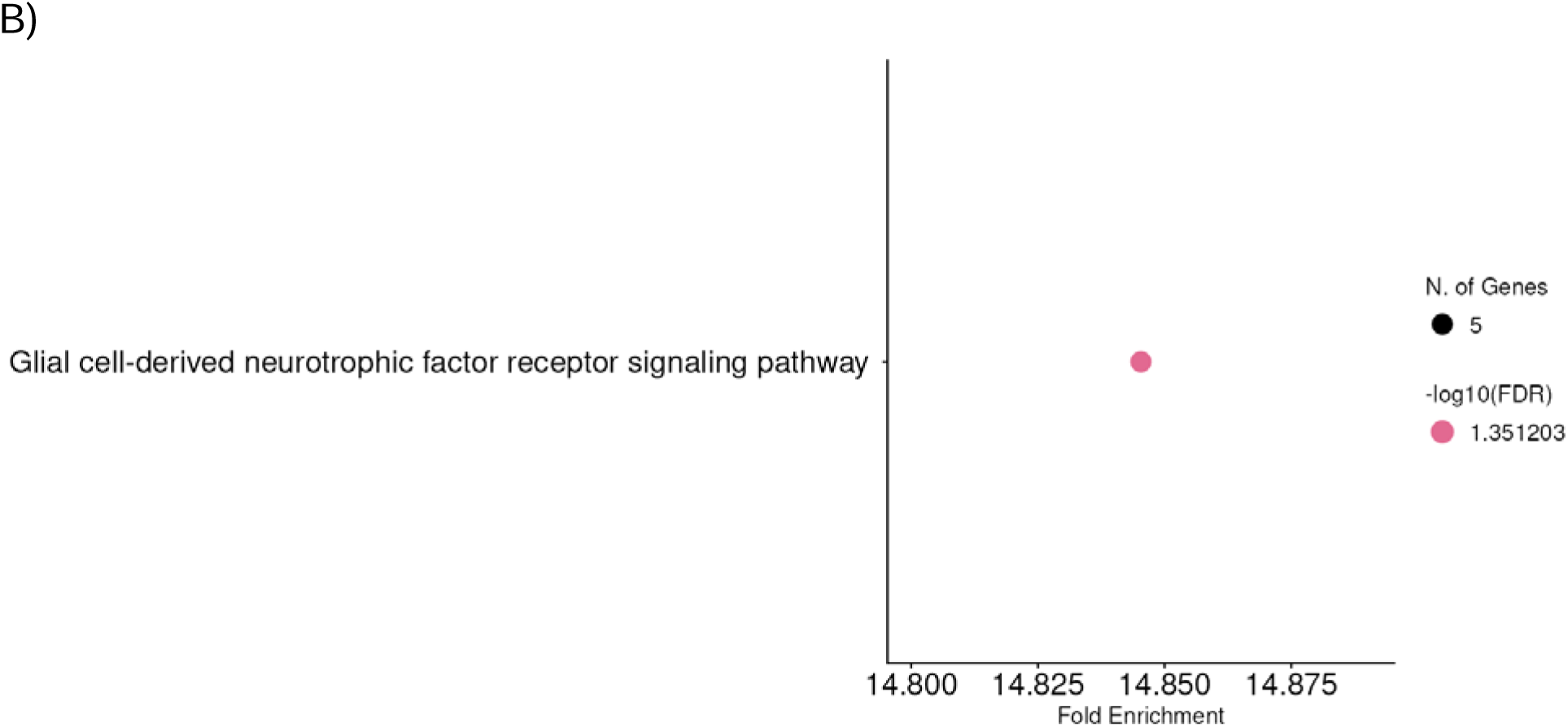
Pathway Enrichment analysis of GeoMx Level 3 results enriched with loading A) In the compressive region (Control Compression vs Loaded Compression) B) In the tensile region (Control Tension vs Loaded Tension) (ShinyGO 0.85).

**Supplementary Figure 5:**
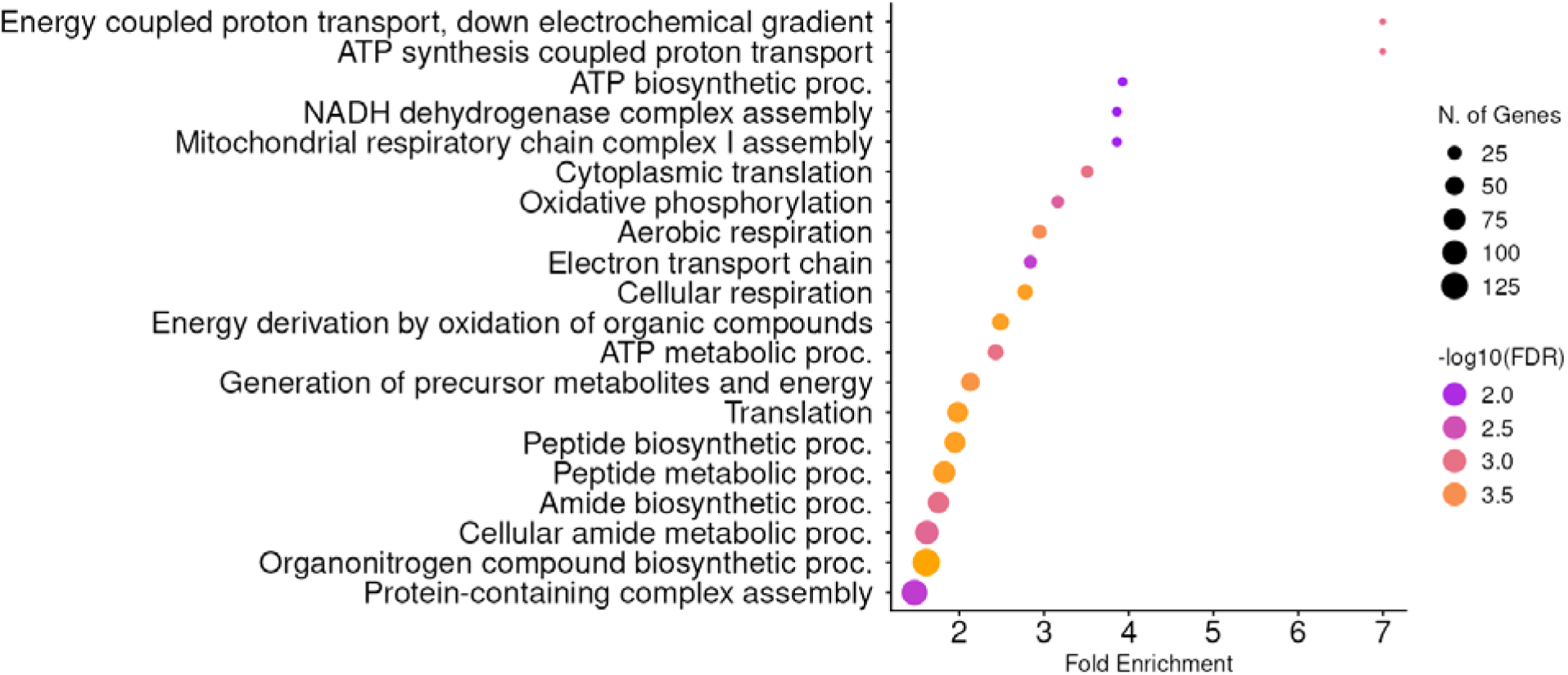
Pathway enrichment analysis of Level 3 GeoMx genes differentially expressed between Loaded Compression and Loaded Tension sites. Only DEGs not significant in the corresponding control comparison (Control Compression vs. Control Tension) were retained to filter out potential contaminants, and GO terms related to muscle were excluded.

**Supplemental Figure 6:**
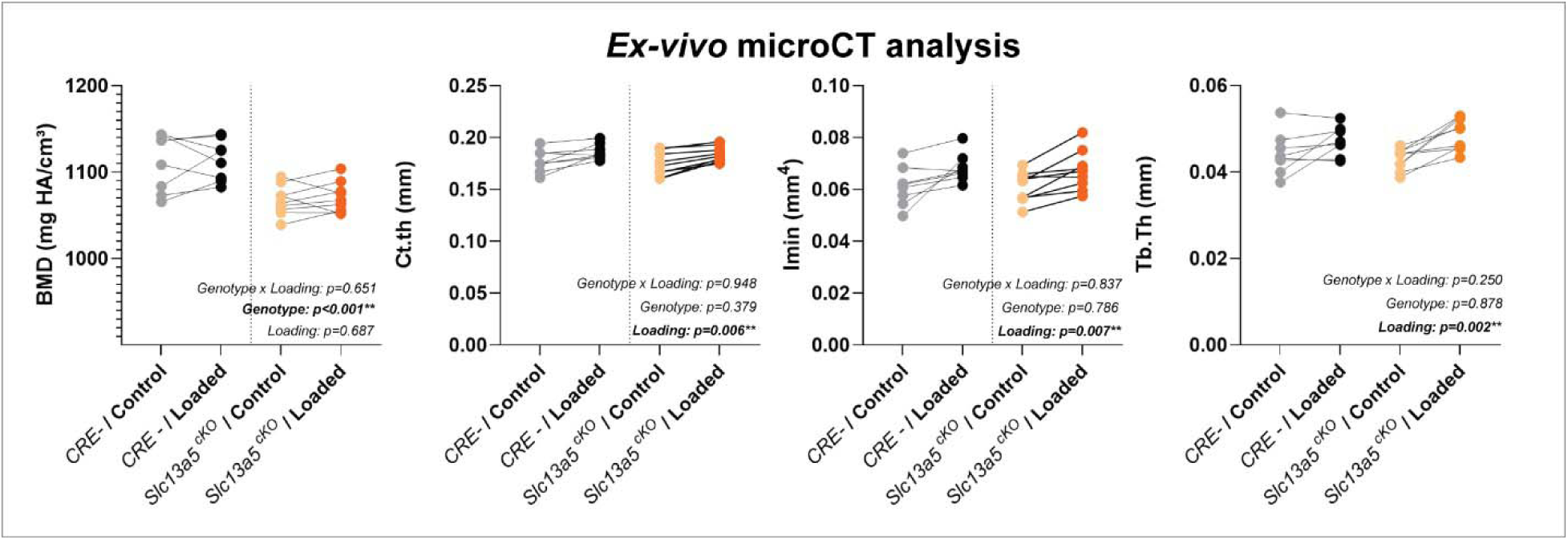
*Ex-vivo* microCT analysis of control and loaded legs from both genotypes. Parameters reported are: BMD, Ct.Th, Imin, and Tb.Th. (*Slc13a5 ^cKo^*: n=9, *Cre- Slc13a5^fl/fl^*n=8). Dots represent each data point, and the line connects the control and loaded tibia from the same mouse. Significant effects of genotype and loading were tested via 2-way ANOVA.

