## Supplemental Results for "Spatial transcriptomics for gene discovery identifies *Slc13a5* as a modulator of bone mechanoadaptation"

**Supplemental Figure 1:** **Pathway Enrichment analysis of GeoMx Level 2 analysis results that are enriched with loading.** A) In the cortical bone (Control Bone vs Loaded Bone) B) In the periosteum (Control Periosteum vs Loaded Periosteum) (ShinyGO 0.85).

**A)**

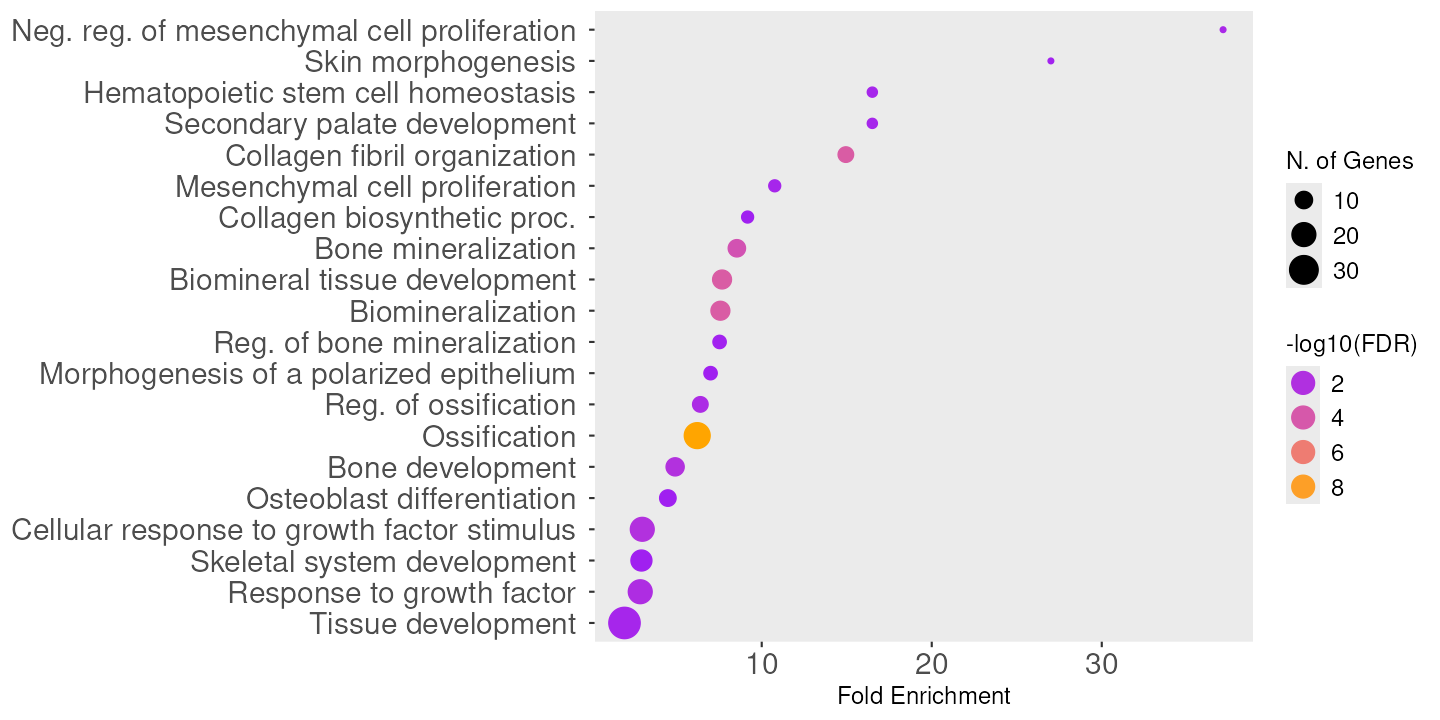

**B)**

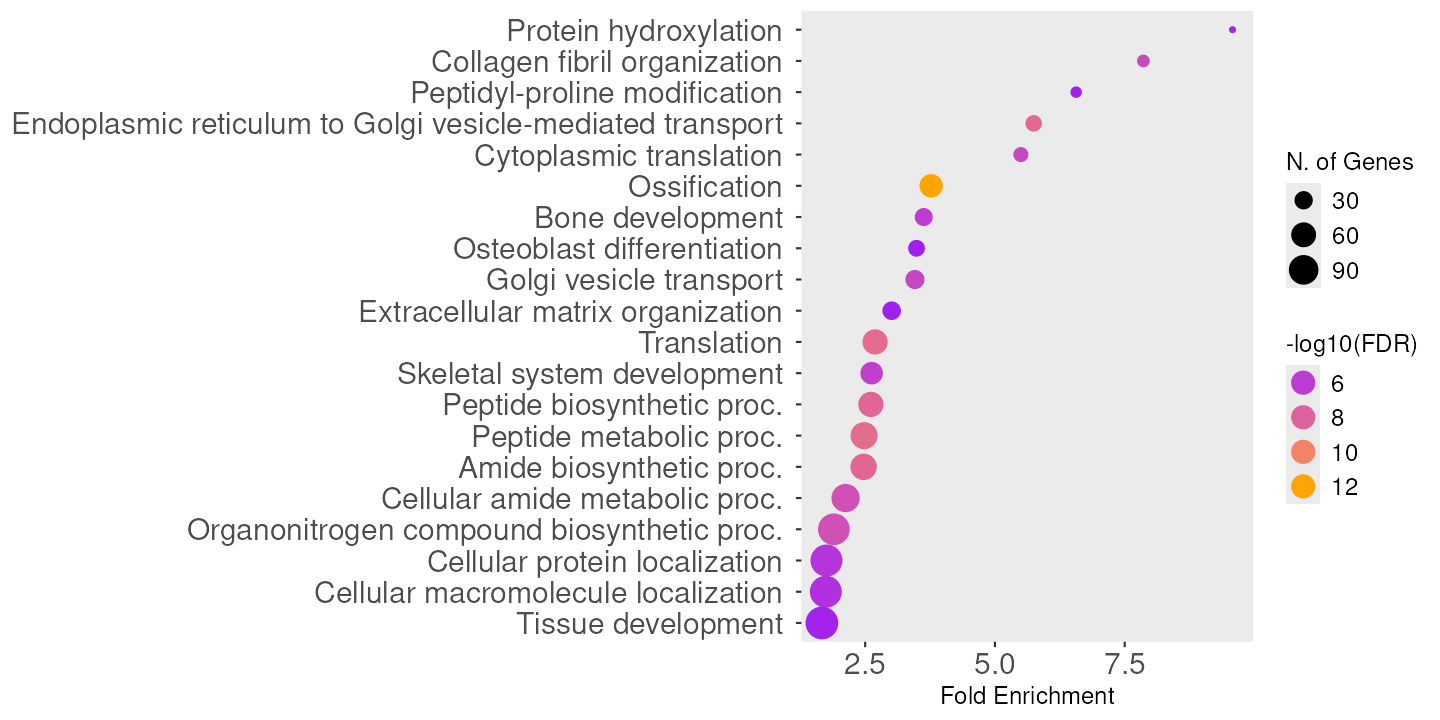

**Supplemental Figure 2: Correlation between GeoMx Level 2 data set results (Control Periosteum vs Loaded Periosteum) and microdissection laser capture data set (Harris et al.)**

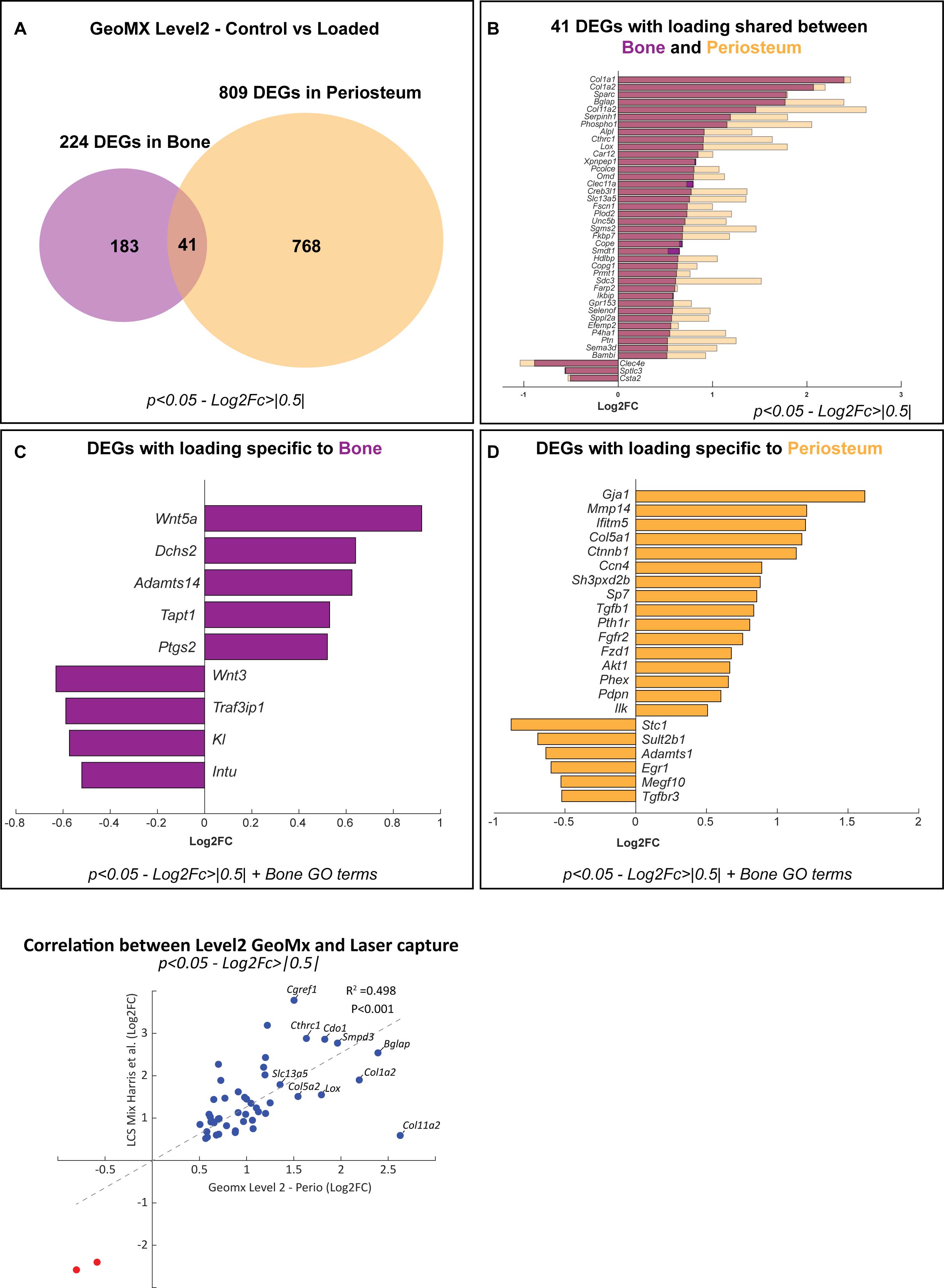

**Supplemental Figure 3: Correlation between GeoMx Level 3 genes enriched in loaded compressive regions (Perio/Bone merged) and genes enriched in loaded tensile regions (Perio/Bone merged).**

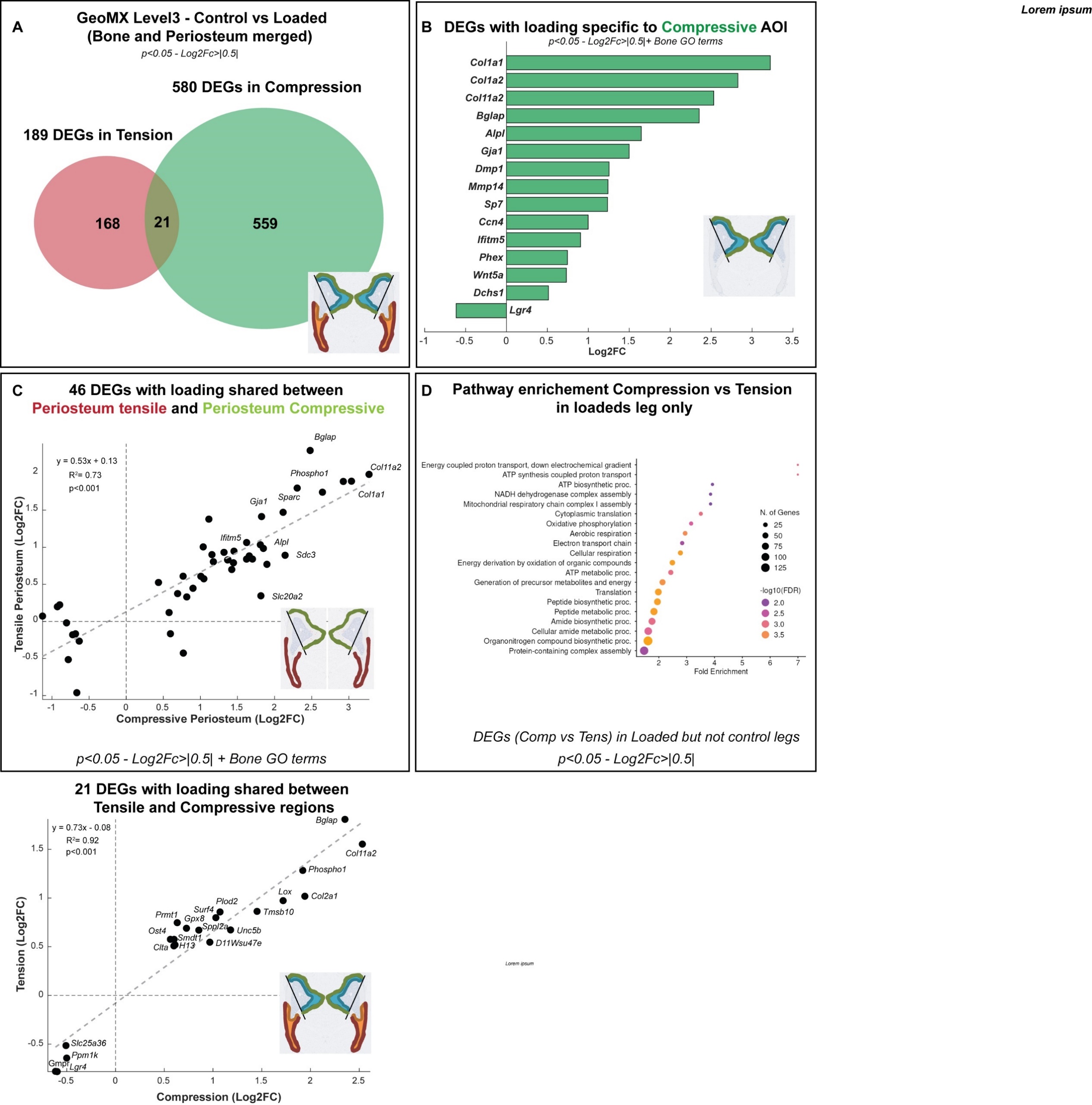

**Supplemental Figure 4:** **Pathway Enrichment analysis of GeoMx Level 3 results enriched with loading.** A) In the compressive region (Control Compression vs Loaded Compression) B) In the tensile region (Control Tension vs Loaded Tension) (ShinyGO 0.85).

**A)**

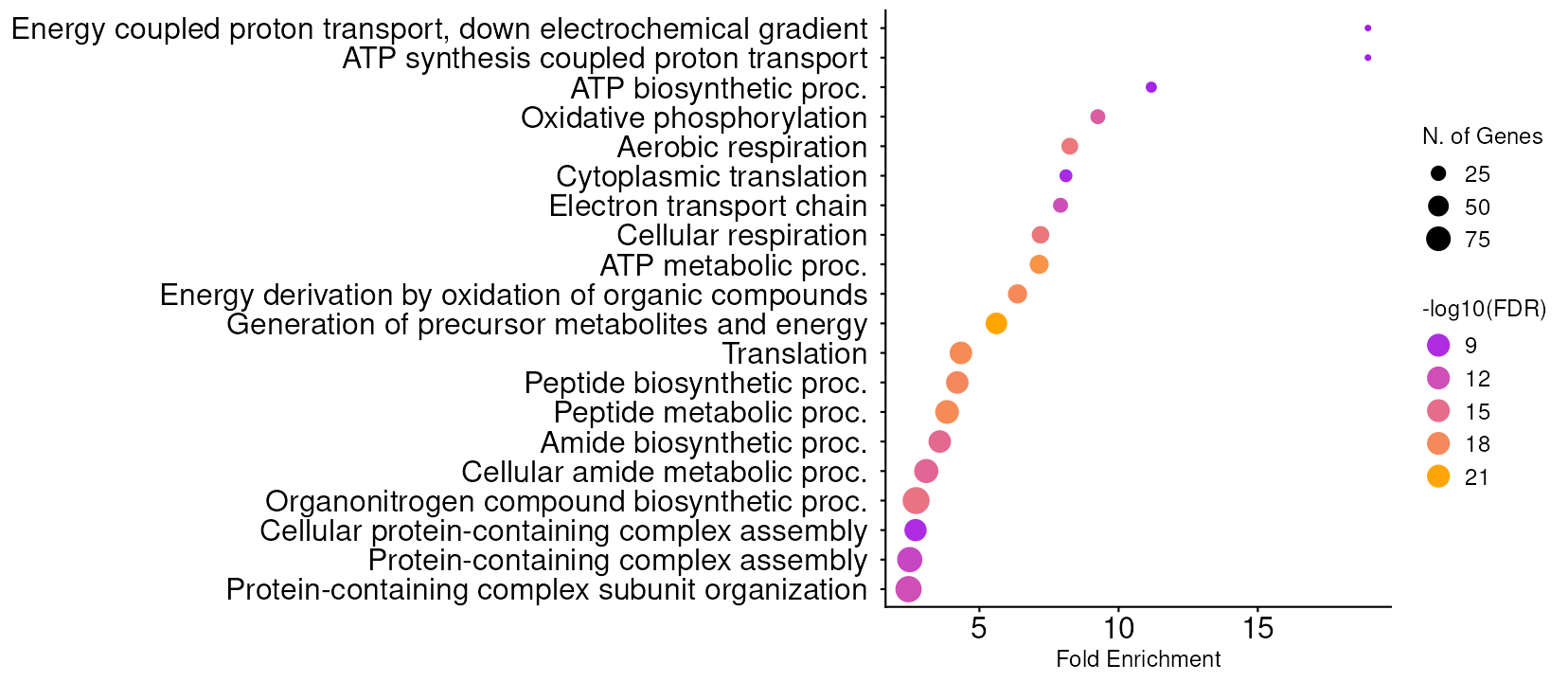

**B)**
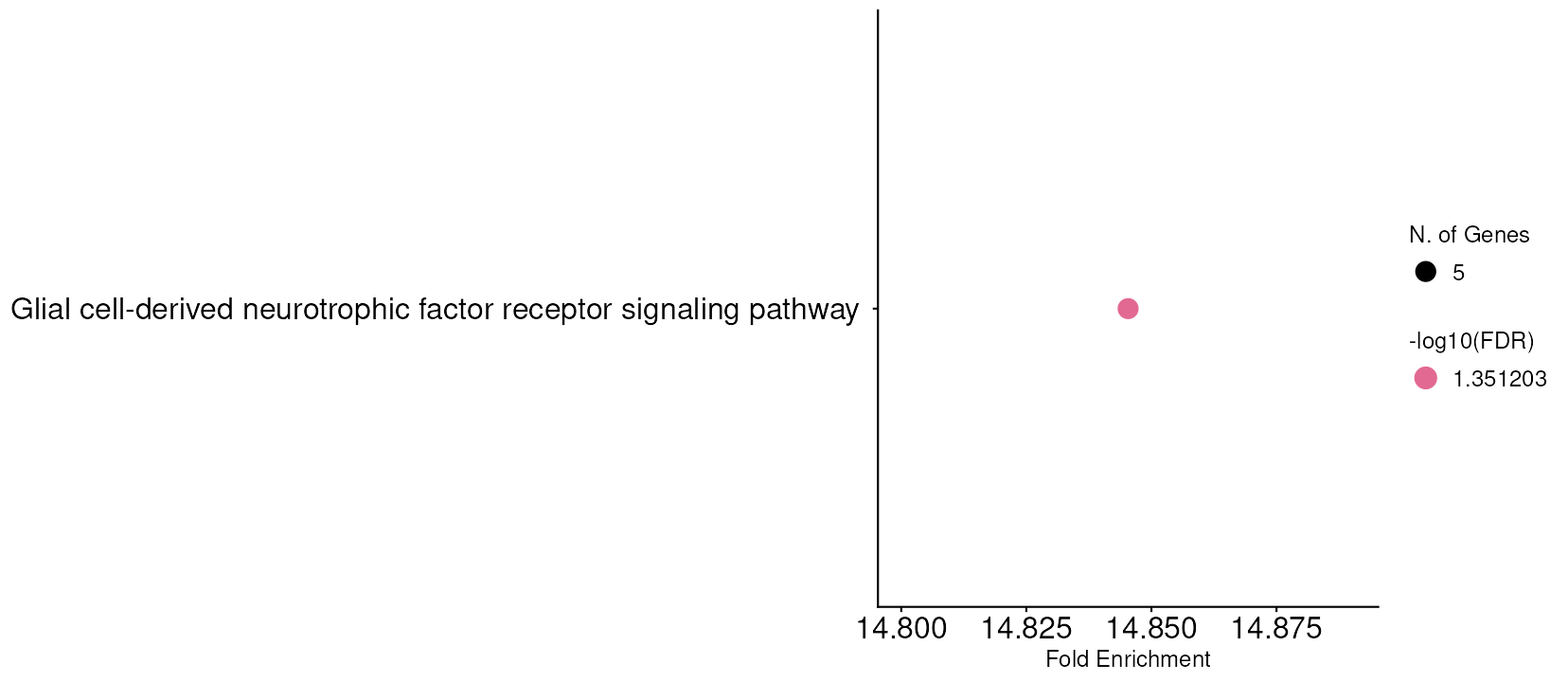

**Supplementary Figure 5: Pathway enrichment analysis of Level 3 GeoMx genes differentially expressed between Loaded Compression and Loaded Tension sites.** Only DEGs not significant in the corresponding control comparison (Control Compression vs. Control Tension) were retained to filter out potential contaminants, and GO terms related to muscle were excluded.

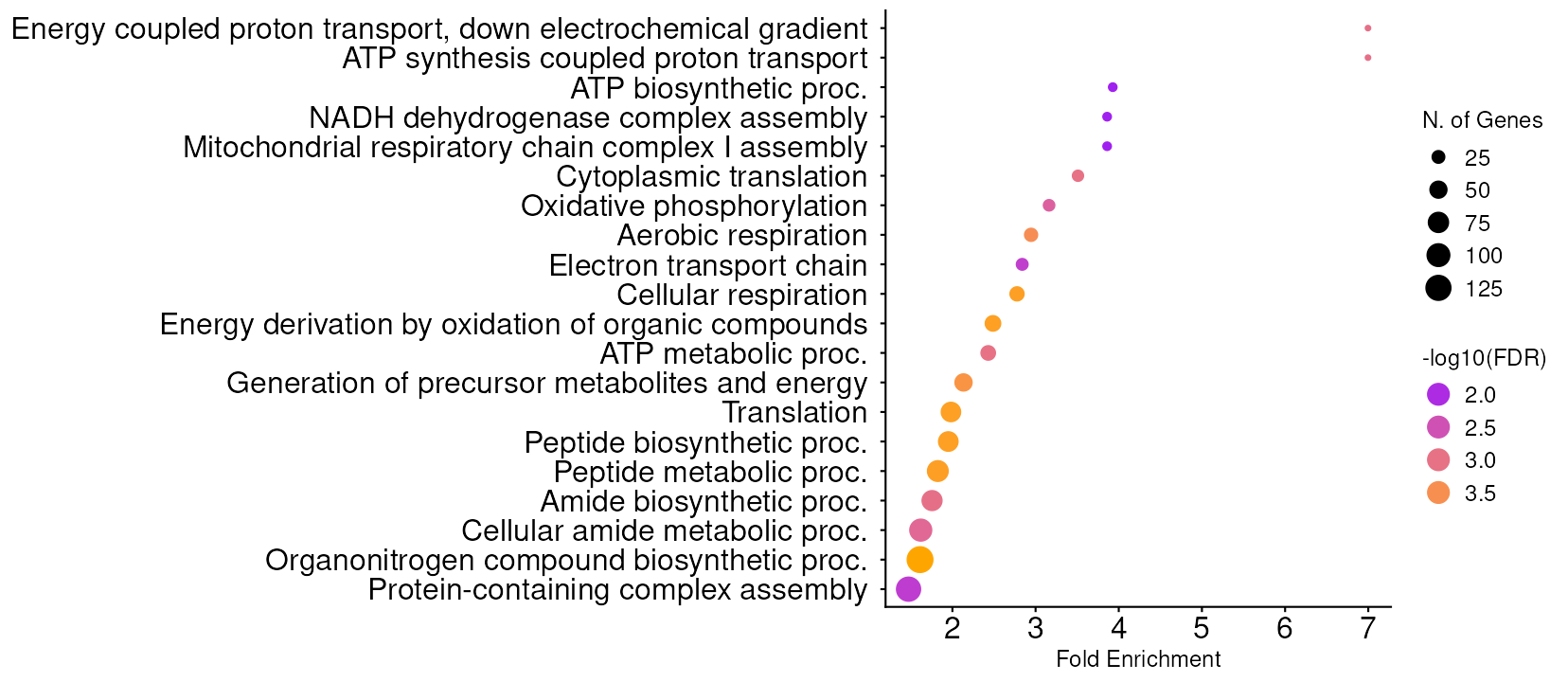

**Supplemental Figure 6: *Ex-vivo* microCT analysis of control and loaded legs from both genotypes.** Parameters reported are: BMD, Ct.Th, Imin, and Tb.Th. (*Slc13a5 ^cKo^*: n=9, *Cre-* *Slc13a5^fl/fl^* n=8). Dots represent each data point, and the line connects the control and loaded tibia from the same mouse. Significant effects of genotype and loading were tested via 2-way ANOVA.

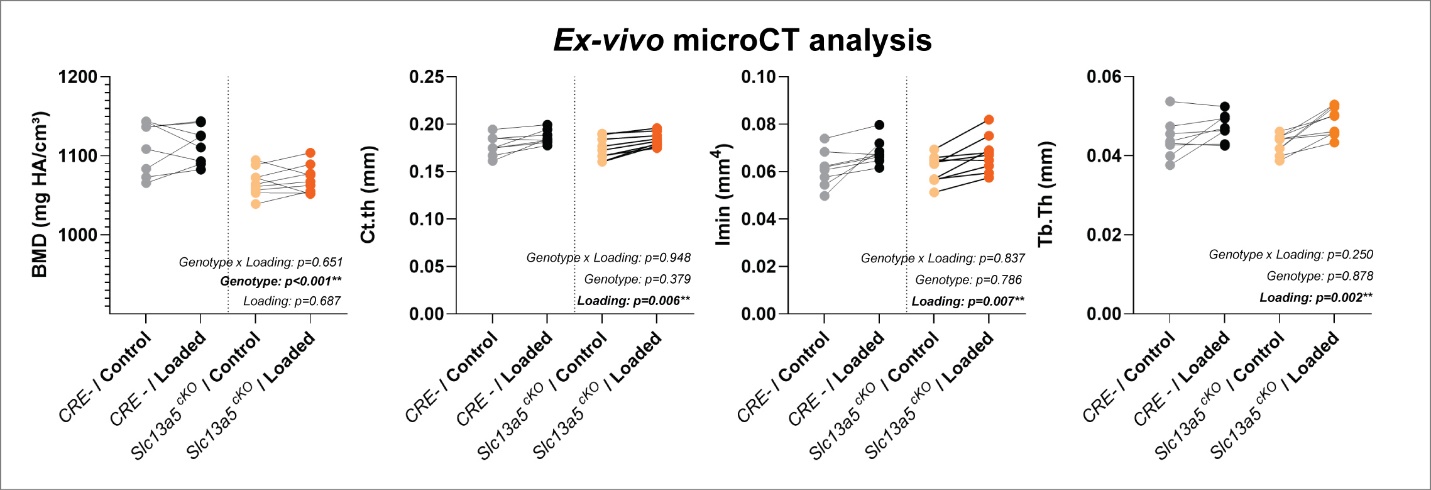
